# A new HIV-1 latency reversing agent activating HIV-Tat

**DOI:** 10.64898/2026.08.20.745916

**Authors:** Phuoc Bao Viet Tong, Laetitia Marty, Nawel Chekrit, Martine Pugnière, Jean-Marie Péloponèse, Célia Bouhnik, Edouard Tuaillon, Alain Makinson, Laurent Chaloin, Bruno Beaumelle

## Abstract

Despite its efficiency to prevent viral multiplication, antiretroviral therapy does not affect HIV-1 latently-infected cells. These cells do not produce significant amounts of viruses and constitute HIV-1 reservoir. To purge this long-lived viral reservoir, the "shock and kill" strategy relies on the use of latency reversing agents (LRAs) to induce activation of latent cells. All LRAs developed until now target cellular proteins and are therefore not specific for HIV-infected cells. Here we present a new LRA that binds and activates HIV-1 Tat which is the key regulator for viral transcription and latency reversal. This molecule termed D10 was designed to bind to the major groove of the Tat protein, and found to activate Tat transcriptional activity by stabilizing the HIV transcription complex. This LRA induces strong HIV production by latent cell lines and latent cells from people living with HIV-1. On latent cells from PBMCs, D10 is active at ∼50 nM, the concentration required to stabilize HIV transcription complex. D10 is the first Tat activator available and the first LRA that targets an HIV protein.

## INTRODUCTION

Antiretroviral therapy (ART) is extremely efficient for suppressing HIV-1 multiplication, but it does not affect the latent HIV-1 reservoir that is established very quickly upon initial infection(Shan *et al*, 2017; Whitney *et al*, 2014). In these latent cells, viral DNA is integrated within the host-cell genome but no significant amount of virus is produced. This is because the transcription (Einkauf *et al*, 2022) and translation (Dube *et al*, 2023; Wu *et al*, 2023) of viral genes is very low in latently infected cells, making them insensitive to ART and difficult to distinguish from uninfected cells by the immune system. HIV reservoir is made of different cell types, essentially quiescent T-cells (Cohn *et al*, 2020), but also monocytes (Veenhuis *et al*, 2023), macrophages (Ganor *et al*, 2019) and microglial cells (Abreu *et al*, 2019). Upon ART interruption, as few as one latently infected CD4+ T-cell that undergoes a stochastic reactivation will be sufficient to cause viral rebound (Hill *et al*, 2014). Hence, even if latent cells are rare, they represent a major obstacle to viral eradication (Sengupta & Siliciano, 2018; Spivak & Planelles, 2018). A strategy to eliminate latent cells is the "shock and kill" approach. In the shock phase of this strategy, latent cells are activated by a latency reversing agent (LRA) to induce viral production, thereby changing the cell phenotype from latent to productive. These infected cells can then be eliminated in the "kill" phase by viral cytopathic effects and/or cytotoxic cells (Board *et al*, 2021; Kim *et al*, 2022). A successful reservoir elimination by powerful LRAs should, together with ART, allow HIV cure while lifelong antiviral therapy is the only available therapeutic strategy at the moment (Rodari *et al*, 2021).

A limited number of molecules have been reported to act as LRAs. Until now, they all target cellular proteins such as histone deacetylase (HDAC), protein kinase C (PKC) or NF-κB (Nixon *et al*, 2020; Rodari *et al*., 2021). PKC agonists such as bryostatin-1 (BST-1) are considered to be the most powerful LRAs (Rodari *et al*., 2021). Nevertheless, both BST-1 and HDAC inhibitors inhibit CD8+ T-cell cytotoxic function *in vitro*. These side effects indicate that the clinical use of these molecules might be difficult due to largely unavoidable off-target cellular effects (Spivak & Planelles, 2018). Hence, there is an urgent need for more specific LRAs (Rodari *et al*., 2021).

HIV-1 Tat is a key regulator of HIV-1 multiplication that positively regulates the expression of viral genes, at the level of transcription elongation especially (Ott *et al*, 2011). Tat is strictly required for the efficient transcription of viral genes from HIV-1 promoter LTR. To this end, Tat binds to the transactivation response (TAR) element on the nascent RNA and cooperatively recruits Cyclin T1 (CycT1) that, together with cyclin-dependent kinase 9 (CDK9), makes up the positive transcription elongation factor b (P-TEFb). CDK9 hyper-phosphorylates the C-terminal domain of RNA polymerase II (PolII), enabling PolII to become processive and produce full-length HIV-1 transcripts (Ott *et al*., 2011). In agreement with its central role in HIV-1 transcription, Tat level and activity were found to be key determinants of latency (Besnard *et al*, 2016; Cary *et al*, 2016; Razooky *et al*, 2015), and Tat induction was found to reactivate HIV-1 latent clones (Razooky *et al*., 2015).

In this study, we describe a Tat activator, termed D10, which is a powerful LRA. This new LRA acts by stabilizing HIV transcription complex Tat-TAR-P-TEFb to which D10 binds with nanomolar affinity. When tested on latent cell lines, and *ex vivo* on latent cells from people living with HIV-1 (PLWH), D10 was at least as efficient as the best LRAs available to date.

## RESULTS

### D10 identification

We aimed to identify Tat-binding compounds that could induce a transcriptionally-hyperactive conformation. A key issue with the identification of Tat ligands using chemoinformatic approaches is that Tat is an intrinsically disordered protein that is poorly folded (To *et al*, 2016). Accordingly, only NMR structures are available for uncomplexed Tat (Bayer *et al*, 1995; Shortridge *et al*, 2018). We performed a molecular dynamics simulation to explore the conformational space used by Tat and identify the most stable Tat conformations. We selected the three conformations of lowest energy (Fig.1A) and tested the ability of compounds from a 55,000 molecules library to fit into Tat main groove. We chose to center the virtual screening on Trp11 because it is somehow buried, so that drug binding was unlikely to perturb Tat interaction with its partners (Bayer *et al*., 1995; Yezid *et al*, 2009). It is also well conserved among Tat from different viral isolates (Debaisieux *et al*, 2012). In agreement with the hydrophobicity of this target pocket, the 30 drugs displaying the highest theoretical affinity (docking scores) were polycyclic compounds predicted to be poorly or not water-soluble. A high degree of structure similarity was observed between hit candidates. They also have a rather elongated shape allowing them to lie within the Tat protein groove (Fig.1B). From the best-ranked compounds, we selected 10 of them according to their diversity of molecular structure (Fig.S1) and tested their cytotoxicity and ability to potentiate Tat transcriptional efficacy using transactivation assays in HeLa cells. These molecules did not display significant cytotoxicity below 10 µM on HeLa cells (Fig.S2A).

**Fig. 1:**
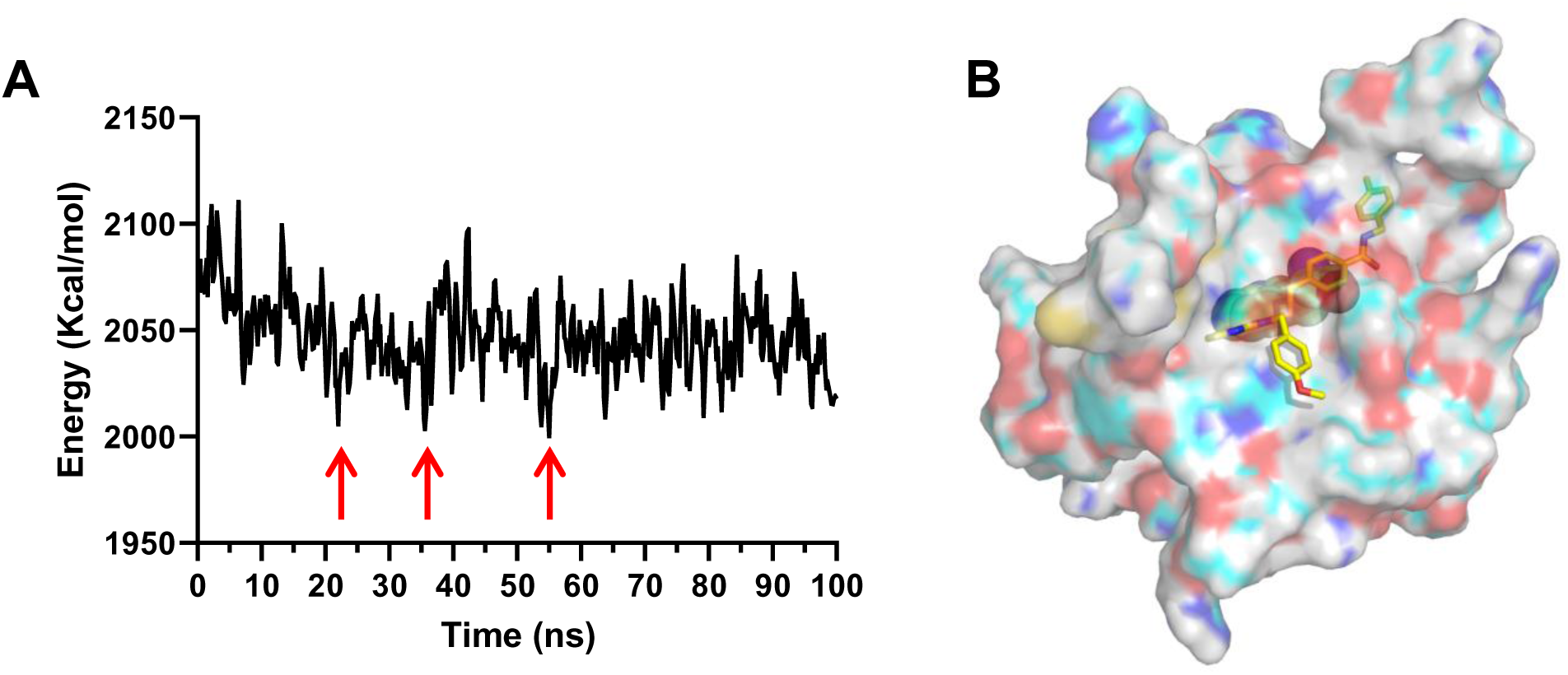
Molecular Dynamic simulations and virtual screening. **A**, Potential energy graph identifying three major stable conformations (*i.e.* conformers with lowest conformational energy) of Tat during 100 ns. They are indicated by red arrows. **B**, Space-filling representation of D10 interaction with Tat obtained by molecular docking. D10 is depicted as yellow sticks and Trp11 side-chain atoms as green van der Waals spheres.

### D10 stimulates Transcription from the LTR in the presence of Tat

Among the Tat-targeting drugs, D10 was found to strongly stimulate transactivation from the LTR (Fig.S2B). We then compared D10 efficiency with other LRAs, SAHA (Vorinostat), an HDAC inhibitor, AZD5582, an NF-κB activator and bryostatin-1 (BST-1), a PKC modulator (Laird *et al*, 2015; Nixon *et al*., 2020; Rodari *et al*., 2021). For these transactivation assays, Tat was either transfected (Tat in), added as a recombinant protein in the medium so that it can enter cells and reach the nucleus (Vendeville *et al*, 2004) (Tat out) or absent (no Tat). D10 facilitated transcription from the LTR only when Tat was present (Fig.2), indicating that D10 selectively stimulates transactivation by Tat. In this assay, D10 efficiency to enhance Tat-mediated transcription was similar to that of AZD5582 and BST-1 that are powerful LRAs but, unlike D10, act independently of the presence of Tat (Fig.2). SAHA was inefficient in this assay because it stimulated transcription from the control TK promoter more efficiently than transcription from the LTR (Fig.S3).

**Figure 2.**
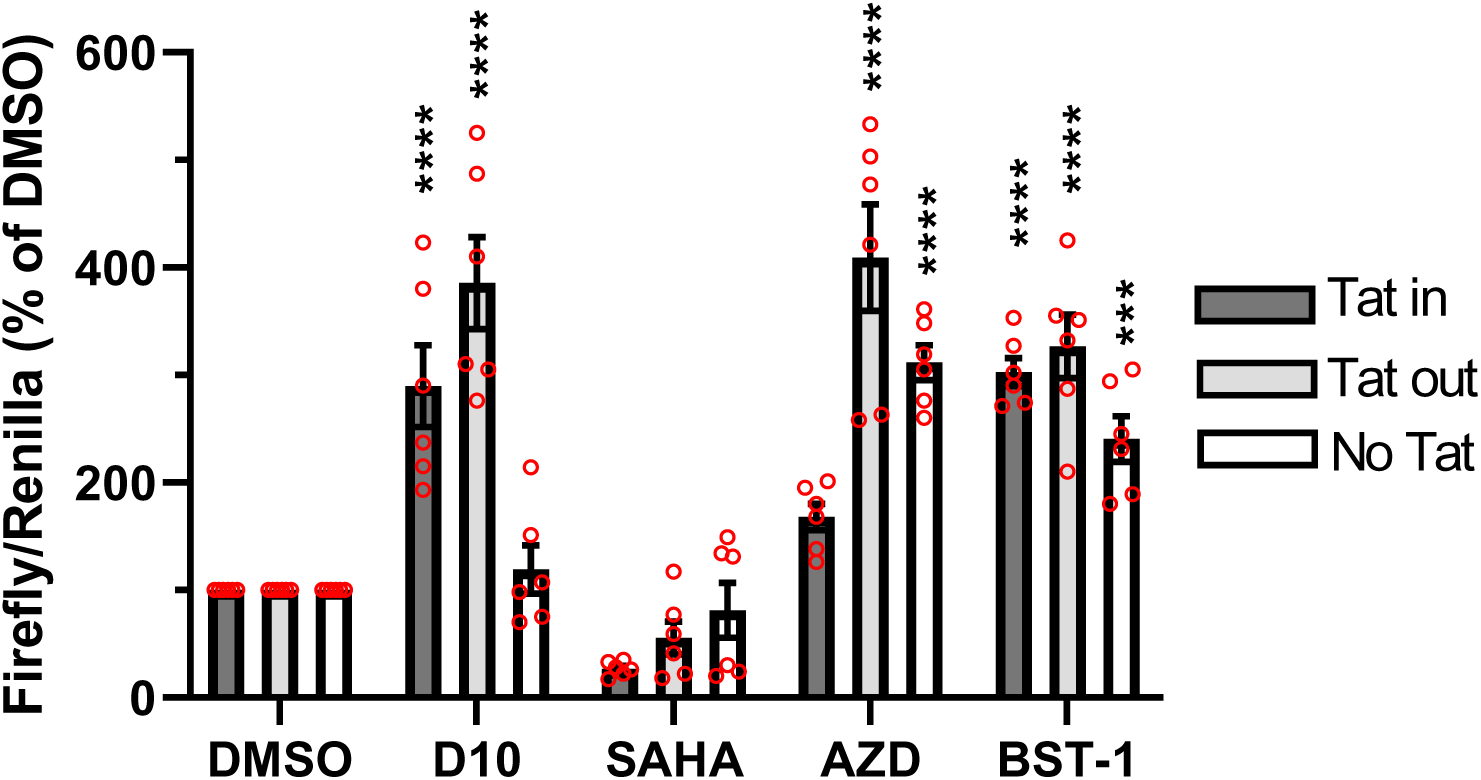
Effect of different LRAs on Tat Transactivation. HeLa cells were co-transfected with a LTR-driven *firefly* luciferase, a *renilla* luciferase behind a cellular thymidine kinase promoter (control vector) and, when indicated ("Tat in"), a Tat expression vector. For the "Tat out" experiments, cells were not transfected with Tat but treated with recombinant Tat (200 nM) 24 h before harvesting cells. No Tat was added in or out for the "No Tat" condition. LRAs (5 µM for D10, SAHA and AZD5582, and 10 nM for BST-1) were added 18 h after transfection for 24 h. Cells were then lysed for luciferase assays. Data are mean ± SEM (n=5-9 independent experiments) and expressed as percentage of solvent. **, p<0.01; ***, p<0.001; ****, p<0.0001 compared to solvent (Two-Way ANOVA).

### D10 is an efficient LRA on cells lines and primary latent cells

We then examined the capacity of D10 to act as LRA on latent cell lines. We first use J-Lat 9.2. This derivative of Jurkat cells harbors a single latent HIV genome in which the Env gene has been inactivated and Nef replaced by EGFP, whose production can be monitored by FACS (Symons *et al*, 2017). D10 LRA activity on this cell line was significant and stronger than that of AZD5582 and BST-1 but lower than that of SAHA (Fig.3). We then used the OM10.1 promyelocytic latent cell line that produces infectious viruses upon induction (Darcis *et al*, 2015). D10 was almost as efficient as SAHA on OM10.1 cells, while BST-1 was the most efficient LRA on this cell line. Such differences in LRA efficiency depending on cell lines were observed before and attributed to differences in cell types and viral genetic background (Ait-Ammar *et al*, 2019).

**Figure 3.**
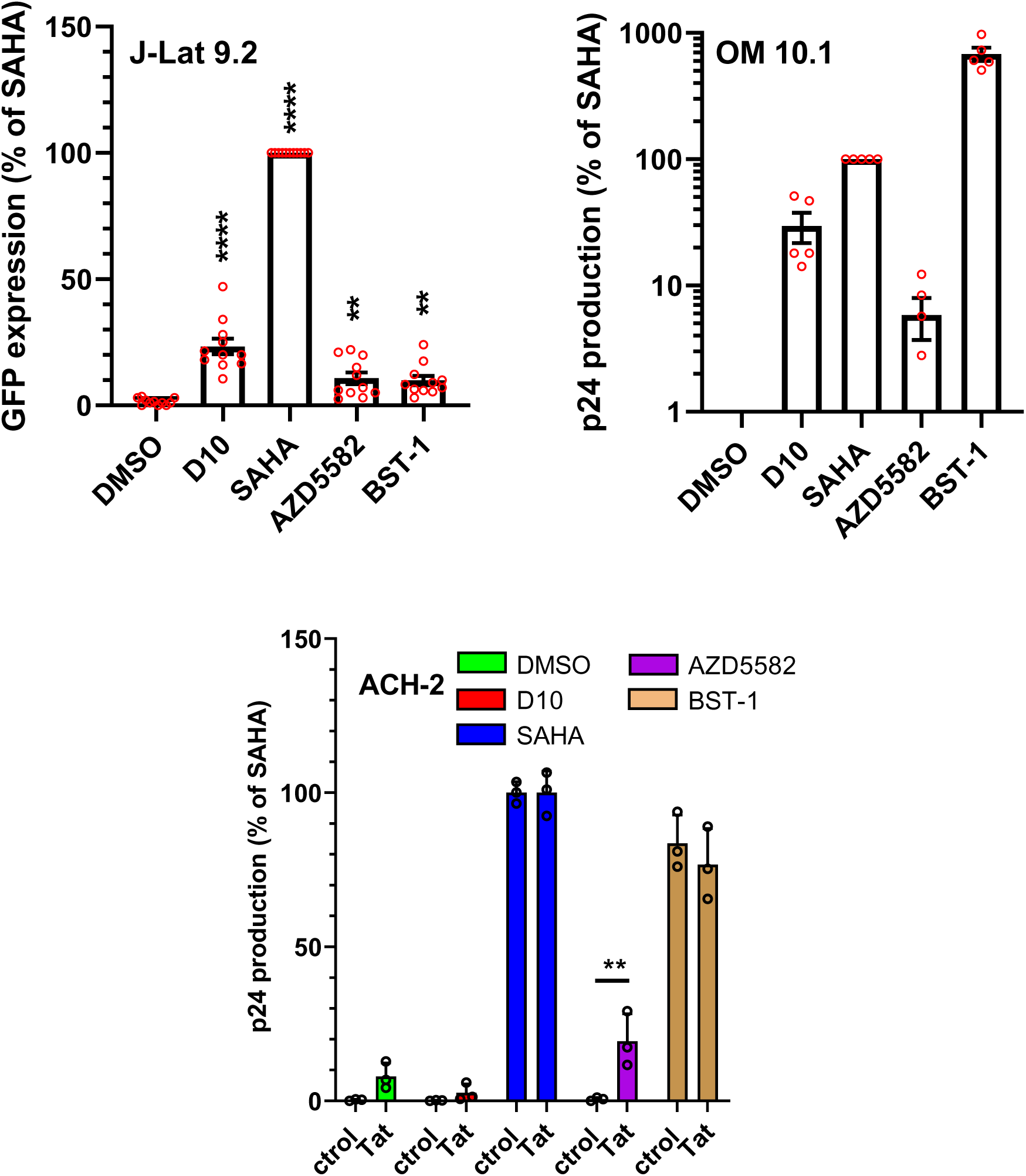
LRA activity on latent cell lines. LRAs (5 µM for D10, SAHA and AZD5582, and 10 nM for BST-1) were added to cell lines for 24 h before harvesting J-Lat9.2 cells for FACS analysis to quantify GFP expression. For ACH-2 cells, 15 nM recombinant Tat was added with the drugs when indicated. The supernatants of OM10.1 and ACH-2 cells were collected after 24h for p24 ELISA. All data are mean ± SEM (n=3-7 independent experiments) and were normalized to SAHA effect that was set at 100%. **, p<0.01; ***, p<0.001; ****, p<0.0001 compared to solvent (One-Way ANOVA for J-Lat 9.2 and OM10.1, and Two-Way ANOVA for ACH-2).

The ACH-2 latent T-cell line has two copies of a latent HIV genome with a defective TAR (Symons *et al*., 2017). This cell line is thus insensitive to the presence of Tat (Cannon *et al*, 1994). When these cells were treated with SAHA or BST-1, a strong viral production was observed, while D10 was unable to induce any viral production by ACH-2 cells, even if exogeneous Tat was added (Fig.3).

These results indicated that D10 LRA activity relies on Tat-TAR interaction, and confirm transcriptional data showing that D10 facilitates transcription from the LTR in the presence of Tat only (Fig.2). It is well established that latent cell lines express minute levels of HIV-1 proteins (Symons *et al*., 2017), and we could accordingly detect significant levels of Tat in J-Lat9.2, OM10.1 and ACH-2 cells (Fig.S4). This expression of Tat likely enables D10 to act as an LRA on the J-Lat9.2 and OM10.1 cell lines. Indeed, as discussed elsewhere, only low amounts of Tat are necessary to insure robust transcription from the LTR (Schatz *et al*, 2023).

Although latent HIV-1 infected cell lines are useful laboratory tools they are clearly different from patient latent cells since cell lines proliferate indefinitely (Xing & Siliciano, 2013). To examine D10 LRA activity on latent primary cells from PLWH, and since latent HIV is present not only in T-cells but also in monocytes (Veenhuis *et al*., 2023), we first used PBMCs depleted of cytotoxic cells. The readout for LRA activity was a p24 ELISA that can detect ∼0.1 pg p24 /ml. To facilitate the comparison between PLWHs, results of these *ex vivo* assays were normalized using BST-1 data.

Results from 18 PLWHs showed that D10 is a robust LRA showing an efficiency of 80-90% compared to BST-1 efficacy. D10 showed a maximum activity within a concentration range of 50-100 nM D10 (Fig. 4A). Similar data were obtained when viral production was monitored using qRT-PCR of vRNA (Fig.S5A), or when purified quiescent CD4^+^ T-cells from the blood of PLWH (Bullen *et al*, 2014; Laird *et al*., 2015) were used (Fig.4B). The LRA activity of D10 on primary latent cells suggests that enough Tat was present in these cells to enable D10 positive effect on Tat transcription. In line with this interpretation, we observed that adding exogenous recombinant Tat, that can enter cells to reach their nucleus (Vendeville *et al*., 2004), did not enhance viral production induced by D10 on latent cells from PLWH PBMCs (Fig.S5B and S5C). The observation that D10 is active on both PBMCs and purified quiescent CD4+ T cells indicate that D10 LRA activity is not dependent on cell types.

**Figure 4.**
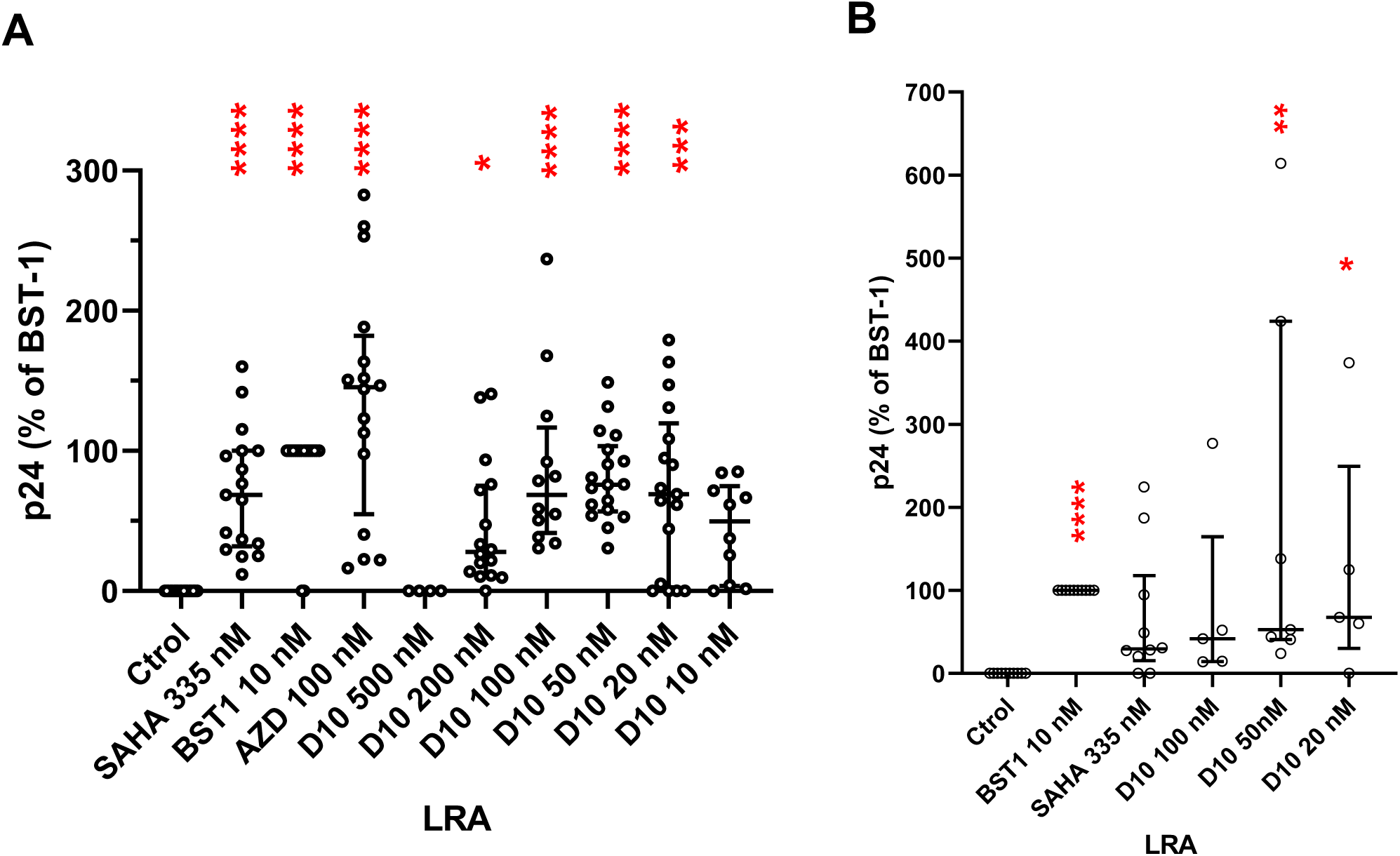
*Ex vivo* LRA activity. **A**, PBMCs from PLWH were depleted from CD8 and NK cytotoxic cells. Cells in duplicated wells were then incubated for 18-20 h in the presence of the indicated concentration of drugs, before collecting supernatants for p24 ELISA. **B,** Quiescent T-CD4 cells were isolated from the blood of PLWH before assaying LRA *ex vivo* activity. Bryostatin-1 (BST1) response was set to 100%. BST1-induced p24 production in the supernatant was within the 0.2-80 pg/ml range, depending on PLWH. Data are median with interquartile range of data from 18 (**A**) or 5-9 (**B**) PLWHs. Kruskal-Wallis tests compared to control; *, p<0.05; **,p<0.01; ***, p<0.001; ****, p<0.0001.

### D10 is not an HDAC inhibitor

D10 owns a benzamide group that is also present in SAHA. We thus examined whether D10 could act as an HDACi. To this end, Jurkat cells were treated for 24 h with D10. SAHA and BST-1 were used as positive and negative controls, respectively. While SAHA induced a strong acetylation of all tested histones (H2A, H2B, H3 and H4), neither BST-1 nor D10 induced a significant acetylation of any histone (Fig.S6). Hence, D10 is not an HDAC inhibitor.

### D10 binds and stabilizes HIV transcription complex

An important question is how can D10 favor Tat transcription (Fig.2). To examine this point, we first performed electrophoretic mobility shift assay (EMSA) using fluorescent-TAR and purified GST-Tat and a protocol adapted from one using radiolabeled TAR (Barboric *et al*, 2000). The Tat-TAR complex was only observed when using WT TAR and GST-Tat and not with a TAR version devoid of the UCU bulge enabling Tat binding, or with GST (Fig.5A). A D10 concentration of 50 nM was enough to enhance Tat binding to TAR by ∼50%. Similar data were obtained using the P-TEFb-Tat-TAR complex (Fig.5B). Hence D10 stabilizes the Tat-TAR complex whether alone or in complex with P-TEFb. Although EMSA gels did not directly measure D10 binding to the transcriptional complex, its stabilization by D10 indicated that D10 binds to the transcriptional complex with a Kd of ∼50 nM. Taking advantage that D10 is fluorescent (Fig.S7A), we monitored D10 interaction with Tat using fluorescence polarization, which is a technique of choice to study protein-ligand interactions (Rossi & Taylor, 2011). We found that D10 binds to Tat with a Kd of ∼ 4 µM, while D10 did not significantly interact with GST that was used as a negative control (Fig.S7b). When D10 binding to immobilized Tat was monitored using surface plasmon resonance, low response values were obtained, as expected for a small molecule (the molecular weight of D10 is 498 Da), and the Kd obtained was again ∼4 µM (Fig.S8). Both biophysical approaches therefore indicated that D10 affinity for Tat is significant but 40 to 80-fold lower than the affinity of D10 for the Tat-TAR complex (Fig.5).

**Figure 5.**
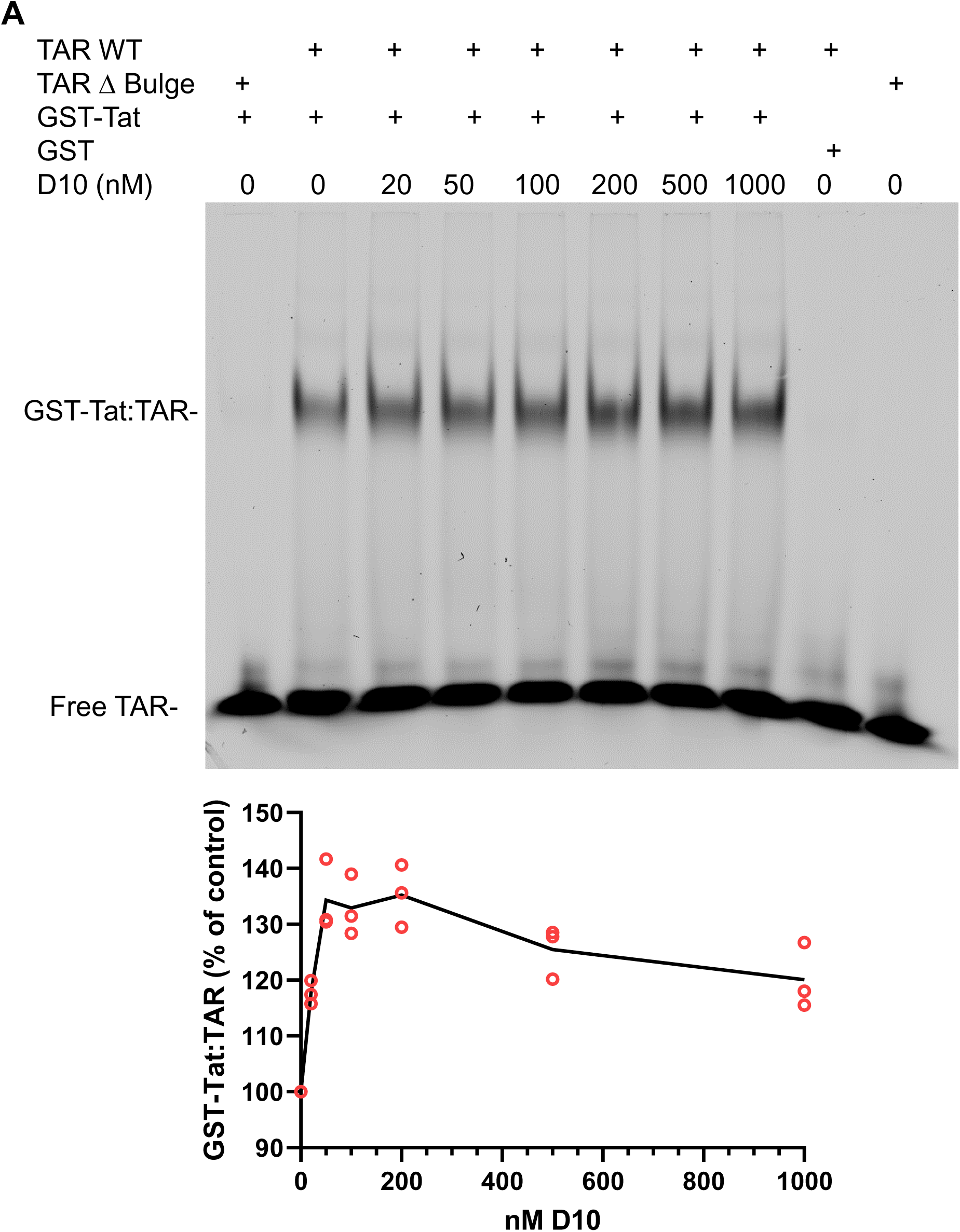
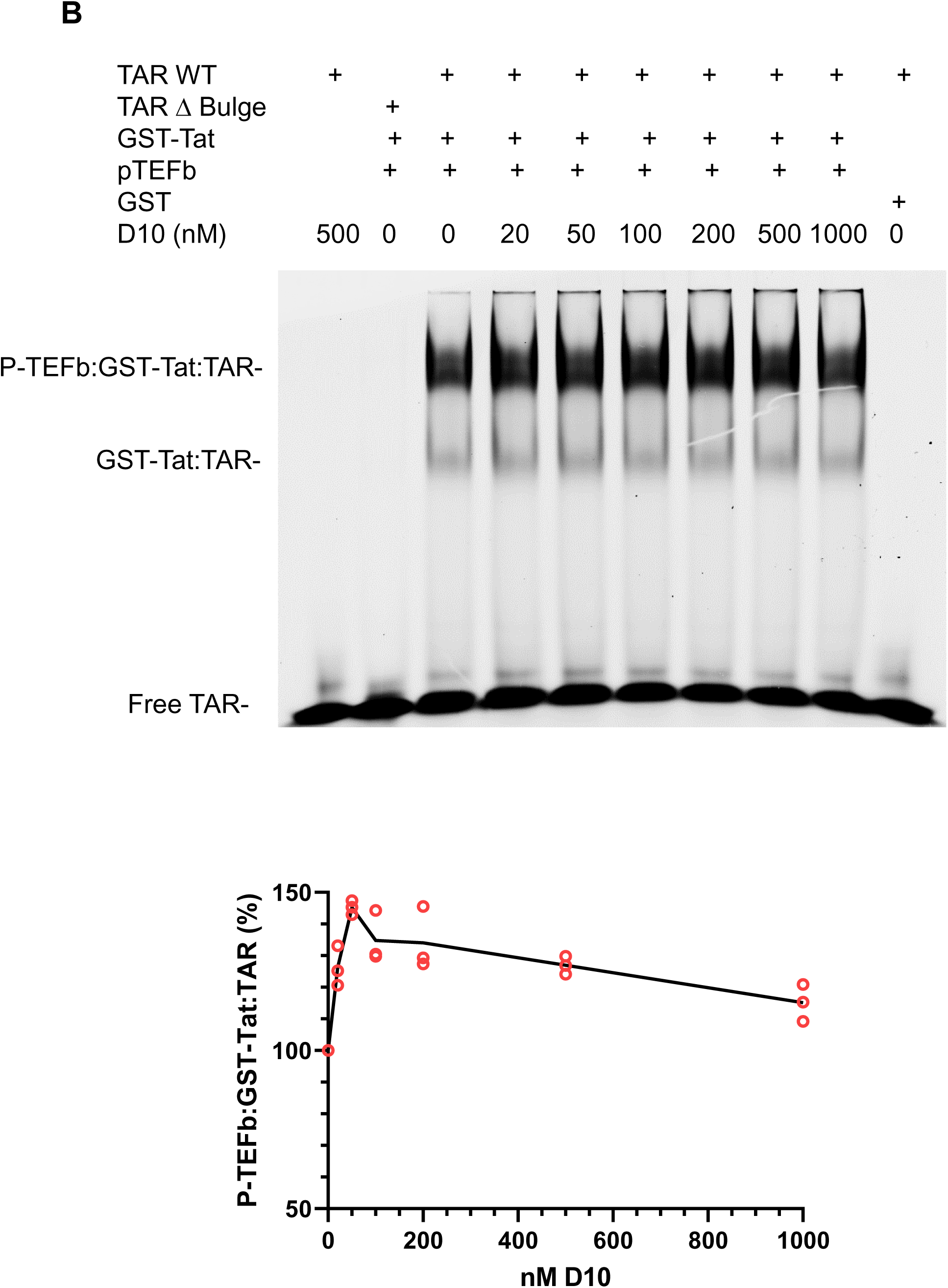
D10 stabilizes the Tat-TAR complex at nanomolar concentrations. **A**, GST-Tat or GST (3 µM) was incubated with 250 nM TAR-FAM either WT or Δbulge for 1 h in the presence of the indicated D10 concentration, before separation on 0.5 X TBE 8% acrylamide gels and FAM fluorescence imaging. The graph shows the quantification from 3 different experiments and the curve connects the means of the triplicates. **B**, the same experiments were performed in the presence of 300 nM purified P-TEFb.

### D10 stabilizes the transcription complex *in cellulo*

We then examined whether D10 could stabilize the transcription complex *in cellulo*. HEK293T cells were transfected with an LTR-luciferase vector, enabling cellular transcription factors such as SP1 and NF-κB to produce TAR from the LTR (Karn & Stoltzfus, 2012). We used nuclear extracts of these cells to pull-down P-TEFb using GST-Tat. The addition of D10 in the nuclear extract doubled the amount of CDK9 and CycT1 recovered after GST-Tat pull down (Fig.6A). This result was confirmed by Tat immunoprecipitation from nuclear extracts of HEK cells. Whether cells were transfected with LTR luciferase or not, *i.e.* whether TAR was present or not, the presence of D10 in cell extracts enabled to double the recovery of CycT1 and CDK9 (Fig.6B). Hence, the strong Tat-CycT1 affinity (∼0.4 nM (Zhang *et al*, 2000)) is sufficient to drive coimmunoprecipitation of the Tat-P-TEFb complex. Collectively, pull down (Fig.6A and 6B) and EMSA (Fig.5) experiments indicated that D10 enables to stabilize HIV transcription complex.

**Figure 6.**
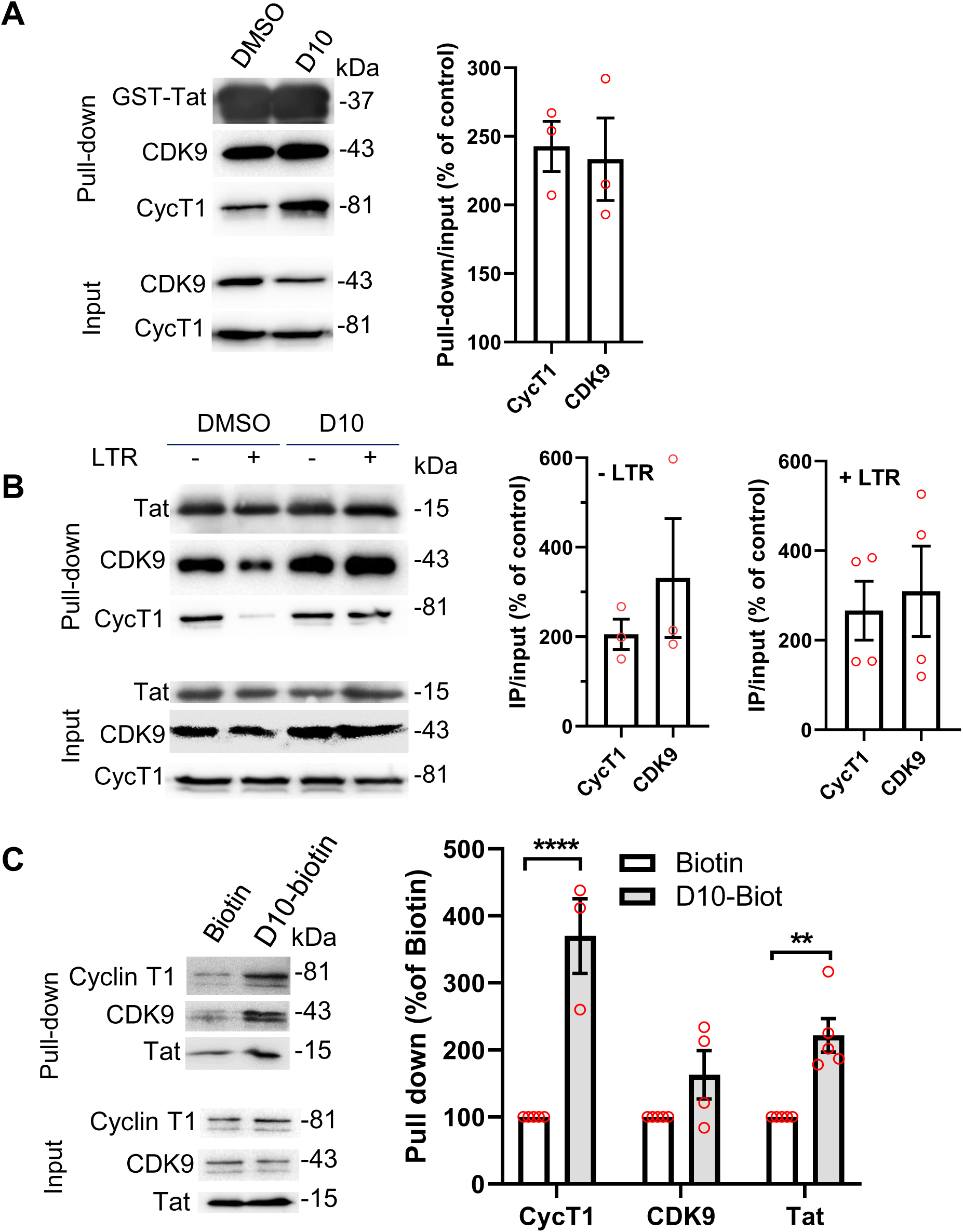
D10 stabilizes HIV transcription complex. **A**, GST-pull down. HEK 293T cells were transfected with pGL3-LTR-luciferase. After 18 h, D10 (5 µM) or DMSO was added for 24 h before preparation of nuclear extracts that received 5 µM D10 when indicated. GST-Tat beads were then added for GST pull down, before SDS/PAGE. **B**, Tat immunoprecipitations. cells were transfected with a Tat-FLAG expression vector with or without pGL3-LTR-luciferase. Nuclear extracts were then prepared, before anti-FLAG immunoprecipitation and SDS/PAGE. **C**, D10-biotin pull-down. HEK 293T cells were transfected with a Tat expression vector and pGL3-LTR-luciferase. D10-biotin or biotin (1 µM) were added to nuclear extracts together with streptavidin magnetic beads before 16 h on a rotating wheel at 4°C, washes and resuspension in SDS PAGE reducing sample buffer. Western blots were stained for CDK9, Cyclin T1 and Tat. Means ± SEM of 3-5 independent experiments. **, p<0.01; ****, p<0.0001 (Two-Way ANOVA).

### D10 binds to HIV transcription complex in cellulo

To examine whether D10 actually binds to HIV transcription complex in cell extracts we used D10-biotin. Biotin was attached using a small linker in place of the Cl atom of D10 (Fig.S1 and S9). D10-biotin enabled to efficiently pull down HIV transcription complex, Tat and CycT1 especially, from nuclear extracts of HEK cells transfected with Tat and LTR-luciferase (Fig.6C). This result indicates that D10 interacts with HIV transcription complex *in cellulo* and that this interaction is strong enough to enable pull down of this complex using D10-biotin.

### D10 enhances the production of long HIV RNAs

Since D10 stabilizes HIV transcription complex it should synergize with Tat to facilitate the production of long RNAs from HIV LTR. We monitored D10 effect on transcription using qRT-PCR and primers enabling to amplify short (∼100 bases), medium (∼5300 bases) or long HIV transcripts (∼8550 bases) (Mousseau *et al*, 2012). Data were normalized using the amount of short transcripts. We noticed that, for HIV-infected Jurkat cells, D10 effect was maximum ∼7 h after adding D10. At this time point, production of medium and long transcripts was enhanced ∼10-fold and 100-fold by D10, respectively (Fig.7A). SAHA, that was used as a control, enhanced the production of medium and long transcripts by 20-30 fold. The enhanced production of long transcripts in the presence of D10 is consistent with an effect on Tat that enhances the production of long transcripts (Ott *et al*., 2011). On HIV-infected Jurkat cells, the effect of D10 and SAHA on transcription was transient and, after 24 h, transcriptional activity returned to normal levels (Fig.7B). This transient effect was less marked when HIV-infected primary T-cells were used, especially regarding the production of long transcripts that was enhanced by ∼4-fold 7 h or 24 h after D10 addition (Fig.7C and 7D). Interestingly, D10 transcriptional effect lasted longer than that of SAHA, which did not show any significant effect on HIV transcription in primary CD4 T-cells after 24h. Altogether these qRT-PCR experiments showed that D10 greatly enhances the production of medium and long transcripts from HIV LTR. D10 effect on HIV transcription is stronger and lasts longer than that of SAHA especially on primary CD4 T-cells.

**Figure 7.**
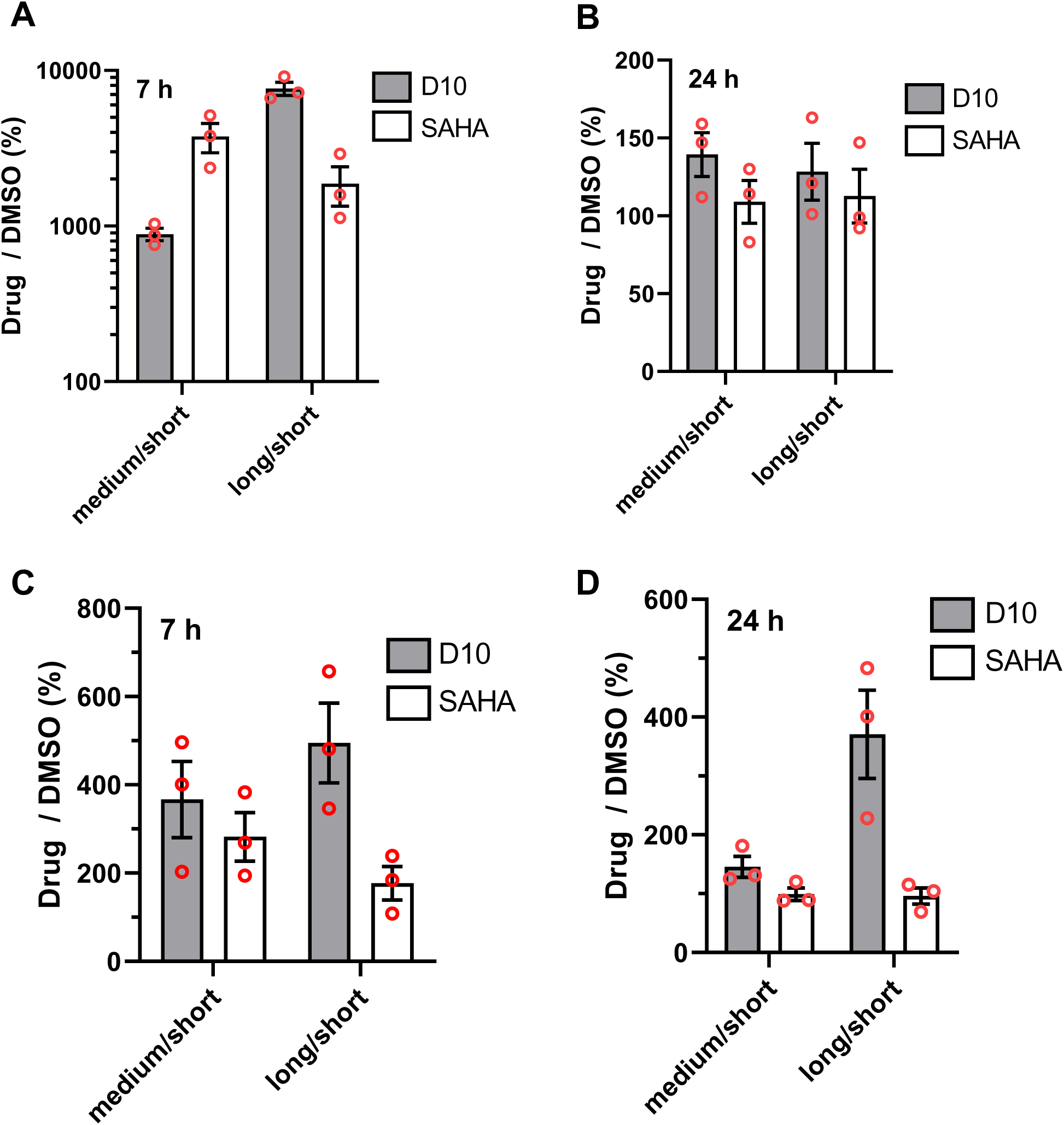
D10 boosts Tat-mediated HIV transcription. **A-B**, Jurkat cells were infected with pseudotyped HIV-1. After 18 h, 5 µM D10 or SAHA were added for 7h (**A)**, or 24h (**B**). Cells were then harvested for RNA extraction, reverse transcription and qPCR using set of primers amplifying short, medium or long HIV RNAs. **C-D**, primary CD4 T-cells were infected with pseudotyped HIV-1. After 18 h, 150 nM D10 or SAHA were added for 7h (**C)**, or 24h (**D**), before harvesting cells for RNA extraction, reverse transcription and qPCR. Results of 3 independent experiments (mean ± SEM) are expressed as percent of control (solvent).

### What is the effective D10 dose?

While D10 LRA effect on latent cell lines requires 5 µM (Fig.3), D10 optimal concentration on primary latent cells in LRA *ex vivo* assays was 20-100 nM (Fig.4). The same difference was observed using Tat transactivation assays: while D10 effect requires to exceed 3 µM on the HeLa cell line (Fig.8A), the optimum D10 concentration on primary T-cells was centered around 30 nM (Fig.8B).

**Figure 8.**
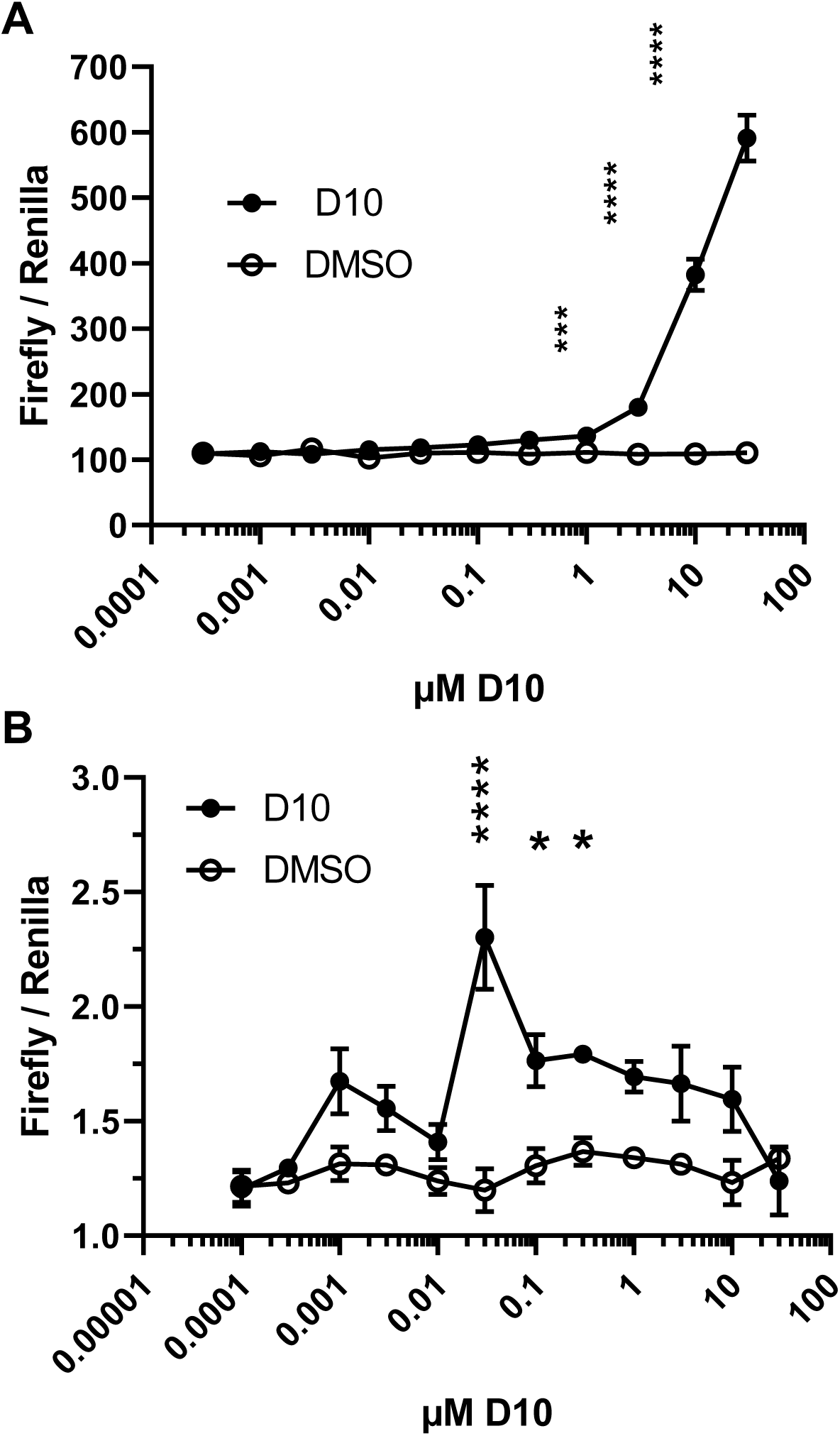
Effective D10 concentrations differ between cell lines and primary T-cells. **A**, HeLa cells were co-transfected with a LTR-driven *firefly* luciferase and a *renilla* luciferase behind a cellular thymidine kinase promoter (control vector) and a Tat expression vector. D10 at the indicated concentration was added 18 h after transfection for 24 h. Cells were then lysed for luciferase assays. **B**, the same experiment was performed using primary CD4 T-cells. Data are mean ± SEM (n=3 independent experiments). *, p<0.05; **, p<0.01; ***; p< 0.001****; p<0.0001 (Two-Way ANOVA).

This hormetic bell-shape response was also observed, albeit less sharply, when we examined D10 ability to stabilize Tat-TAR interaction in EMSA assays (Fig.5). We thought that the requirement of transformed cell lines for high D10 concentration to achieve a biological effect might be due to their capacity to expel drugs using transporters, thereby decreasing D10 intracellular concentration below the active dose. ABC transporters are often overexpressed in cancer cells (Muriithi *et al*, 2020).

Among the different family of ABC transporters, ABCG2 appeared as a strong candidate to export D10 since ABCG2 has been documented to export molecules with physicochemical properties similar to D10, *i.e.* polycyclic and elongated compounds (Mao & Unadkat, 2015). If ABCG2 is responsible for D10 export from cancer cell lines, then ABCG2 inhibitors should enable much lower D10 doses (*i.e* within the nanomolar range) to facilitate Tat transcription in cell lines. To examine this point, we used fumitremorgin C, a specific inhibitor for ABCG2 (Seamon *et al*, 2006). The presence of this ABCG2 inhibitor enables minute D10 concentrations, especially 10 nM, to facilitate transactivation by Tat compared to DMSO (Fig.S10). Cyclosporin A and Psc833 that inhibit other ABC transporters, *i.e.* ABCB1/MDR1 and ABCC1/MRP1 (Cole *et al*, 1994; Engle & Kumar, 2022) did not facilitate D10 effect on Tat transactivation. These results indicated that ABCG2 is responsible for D10 export by cancer cell lines and for the requirement on micromolar D10 concentrations to achieve on cell lines a significant effect that is observed in the nanomolar range on primary cells.

## DISCUSSION

A well-accepted strategy to reduce the size of the HIV-1 reservoir is the shock and kill approach that involves the use of LRA to trigger viral production by latent cells that will thus become producing cells and will be killed by viral infection, or cytotoxic T-cells (Board *et al*., 2021; Kim *et al*., 2022). Available LRAs such as HDACi and PKC modulators can functionally impair the cytotoxic activity of CD8 or NK cells (Spivak & Planelles, 2018). More generally, most LRAs have been designed to manipulate cellular transcriptional or epigenetic regulators, such as HDACs, PKC, or NF-κB. This approach is inherently associated with pleiotropic effects because these pathways control numerous physiological functions. We here describe a new LRA that specifically activates Tat-mediated transcription from the LTR and is a powerful LRA at 20-100 nM *ex vivo* on latent cells from PLWH (Fig.4). This new LRA, D10, is a polycyclic and elongated molecule that was predicted by molecular docking to insert within Tat major groove. This central groove has been observed in Tat structures from several RMN studies (PDB 1TIV, 1TAC and 1TBC (Bayer *et al*., 1995)). Tat residues that interact with D10 are well conserved among the major subtypes B and C (Fig.S11). Interestingly, EMSA (Fig.5) and biophysical experiments (Fig.S7 and S8) indicated that D10 affinity for Tat is enhanced by ∼50-fold when Tat is bound to TAR. Hence, D10 essentially targets Tat when present in the transcription complex.

The absence of viral production induced by D10 on ACH-2 cells that have a latent HIV-1 genome with a TAR unable to bind Tat (Fig.3) confirmed that D10 requires the presence of Tat to act as an LRA. Hence, the levels of Tat expressed by latent cell lines such as J-Lat9.2 and OM10.1 (Fig.S4) is sufficient for D10 to trigger significant viral production by these cells (Fig.3). Regarding primary cells, the LRA activity of D10 on latent cells from PLWH (Fig.4) indicates that these cells contain at least some minute levels of Tat. Very low amounts of Tat are indeed needed to activate transcription (Schatz *et al*., 2023). This Tat can be produced by latent cells themselves since, although most of them contain defective proviruses, viral DNA can be transcribed and translated to some extend (Einkauf *et al*., 2022; Pollack *et al*, 2017). Alternatively, these cells could have internalized circulating Tat produced in *trans* by infected cells (Shmakova *et al*, 2024). The absence of effect of exogenous Tat on D10 LRA activity during *ex vivo* assays (Fig. S5b and S5C) indicated that, whatever its origin, Tat level in these cells is sufficient to sustain D10 facilitating effect on HIV transcription. D10 is the first activator described for Tat transcriptional activity. Its LRA activity indicates that it is possible to obtain LRAs that target HIV proteins, thereby improving LRA specificity.

## MATERIALS AND METHODS

### Molecular Dynamic simulations and virtual screening

Simulations were performed using the program NAMD v2.12 (Phillips *et al*, 2005) in explicit water at constant temperature (300 K) and pressure (1 atm) using the Charmm force-field and c36m topology (Zhu *et al*, 2012). The Tat protein RMN structure PDB 1TIV was chosen because it was obtained at pH 6.5 in contrast to 1JFW, 1K5K or 1TAC that were obtained at pH 4.5-5 where, in our hands, Tat precipitates (Yezid *et al*., 2009). The Tat protein was first immerged into a water box (TIP3P model) then neutralized by the addition of chloride ions and replicated in each direction using periodic boundary conditions. The short-range Lennard-Jones potential energy (Lennard-Jones) was smoothly truncated from 10 to 12 Å and long distance electrostatic was calculated using the Particle Mesh Ewald (PME) algorithm (Essmann *et al*, 1995). The potential energy of the solvated system was minimized for 50 ps using the conjugate gradient algorithm (time step of 2 fs) and the system was progressively heated from 0 to 300 K by steps of 10K. After a 100 ps of equilibration, simulation was performed for 100 ns. Longer simulations (500 ns) have been performed to ensure that we sampled reasonably the conformational space explored by the versatile Tat protein (Fig.S12). Surprisingly, the 500 ns simulation produced a similar structural behavior as observed in the short production run (similar RMSD values for backbone atoms and fluctuations higher for external parts but similar for the core domain). Although Tat has large unstructured parts which may increase its flexibility, no partial protein structuration or folding could be observed within this time scale. Conformational energy (the sum of internal bond and angular energies) was computed from the simulation trajectory in order to identify the most stable protein conformers. Three lowest-energy conformations (Fig.1A) were selected for the remaining of the study and their potential energy was minimized for 50 ps prior docking. All molecular dynamics simulations were analyzed with the VMD software (Humphrey *et al*, 1996).

Virtual screening was performed using the GOLD v5.6 software (CCDC (Jones *et al*, 1997)) on the three lowest-energy conformations identified by molecular dynamics. A spherical area with a 12 Å-radius around Trp-11 was targeted for the search of docking solutions by applying 20 genetic algorithm runs. The different docking orientations of bound molecules were classified according to their docking score calculated by the Goldscore scoring function that evaluates the interaction energy between the screened compound and Tat residues. The different docking poses were analyzed by the clustering method (complete linkage) from the rmsd matrix of ranking solutions. We first screened a subset of the "ChemDiv" (http://www.chemdiv.com/) library that was chosen both because of its small size (55,000 compounds) and its structural diversity. A larger chemical library (including Acros Organics and a subset of drug-like compounds from MolPort) giving a total of 200 000 compounds was then screened. From these two screenings, 30 compounds were selected from the best-ranking hits, reduced to 10 according to their molecular diversity and these D1 to D10 molecules were ordered for biological assays (Fig.S1).

### Chemicals and proteins

All chemicals were of the highest purity available and, except when otherwise indicated, they were obtained from Sigma. Drugs were purchased from MolPort. D10 and D10-biotin were synthesized by AGV discovery (Montpellier, France). Recombinant Tat (BH10 version; 86 residues), GST-Tat and p24 were produced as described (Hung *et al*, 2013; Schatz *et al*., 2023; Vendeville *et al*., 2004). Human P-TEFb (human CDK9/CycT1 (Flag-tagged) was produced and purified from baculovirus infected Sf9 cells as described (Lopez Martinez *et al*, 2025) using the pBIG1e-CDK9/Cyclin T1 from addgene (#231732). Fluorescent TARs (RNA nucleotides 16-43-FAM) were purchased from Sigma. The Δbulge version was devoid of the UCU bulge enabling Tat binding (Ott *et al*., 2011). Anti-CDK9 (CST-2316) and anti-Cyclin T1 (CST-81464) were from Cell Signaling Technology and anti-Tat (sc-65912) from Santa Cruz Biotechnology.

### Cells

Jurkat cells (clone E6-1) were obtained from the ATCC. J-Lat 9.2 is a Jurkat-based latent cell line containing an integrated latent HIV-1 genome that expresses GFP in place of Nef and has a frameshift mutation in Env. J1.1 cells are also latent cells derived from Jurkat cells (Symons *et al*., 2017). ACH-2 is a T cell line with an integrated HIV-1 copy with a TAR mutation preventing Tat binding to TAR (Symons *et al*., 2017), OM10.1 cells are HL60-derived promyelocyte cells that contain a single integrated provirus (Taura *et al*, 2015). ACH-2, OM10.1, J1.1, J-Lat 9.2 and HeLa cells were from the NIH AIDS Reagent Program. Primary CD4-T cells were purified from the blood of healthy donors (provided by the French Blood Establishment under the agreement 21/PLER/MTP/CNR11/2013-049). After isolating PMBCs by centrifugation on Ficoll-Paque, monocytes were depleted by adherence and T-CD4 cells were purified using a negative selection kit (Miltenyi Biotec 130-096-533). They were then activated using phytohemagglutinin (1 μg/ml) for 24 h then interleukin-2 (50 U/ml) for 6 days.

The blood from PLWH people was obtained under the contract RECHMPL23_0276-CNRS 293537 between Montpellier Hospital (CHU) and the IRIM laboratory. Characteristics of PLWH that provided blood for this study (20ml) are presented in Table S1. Cell purification was started less than one hour after blood-drawing on citric acid-dextrose. PBMCs were isolated on Ficoll-Paque. They were then either depleted of CD8^+^ or NK (CD56^+^) cells using Stem Cell kits # 17853 and #17915. Cell purity was monitored by FACS. In pioneer experiments, PBMCs from PLWH were used to isolate CD4-quiescent T-cells. To this end, T-CD4 cells were obtained by negative selection (Miltenyi Biotec 130-096-533), and resting cells (CD25^-^, CD69^-^ and HLA-DR^-^) were isolated by negative selection as described (Bullen *et al*., 2014; Laird *et al*., 2015) using Miltenyi Biotec appropriate kits (130-097-044, 130-092-355, 130-046-101).

### Cytotoxicity assay

Cells were diluted to 0.8 ×10^6^ cells/ml and 0.1 ml was transferred into wells of a 96-well white plate. Drugs at 5 mM in DMSO were diluted in medium and 0.1 ml of diluted drug was added to each well (n=2). After 24 h of incubation at 37°C, the plates were centrifuged at 1500 rpm for 1 min and 100 μl of supernatant was removed to assay cell viability using the Celltiter-Blue reagent (Promega) as described by the manufacturer. After 4 h incubation at 37°C the fluorescence was measured (excitation 560 ± 20 nm and emission 590 ± 10 nm) using a Tecan SPARK 10M.

### Transactivation and histone acetylation assays

HeLa cells were plated (30 000 cells/well) in 96-well plates. After 24h they were transfected using polyethyleneimine max (40K; polysciences) as described (Longo *et al*, 2013) with a vector expressing a Firefly gene under the control of an LTR promoter, a plasmid with a Renilla gene under a CMV promoter (internal control) and a Tat expressing vector at a ratio 1/5/1.

After 18 h of incubation at 37°C, the medium was changed and drugs were added at 5µM for 24h. When indicated (Tat out) the assay was performed using extracellular Tat. In that case, cells were transfected with luciferase vectors only and recombinant Tat (200nM) was added with the drugs (Vendeville *et al*., 2004). Luciferase activity was then assayed as described by the manufacturer (Dual-Glo luciferase assay, Promega). Briefly, the cell medium of each well was replaced by 30 μl of PBS. Then 30 μl of Dual-Glo reagent was added. Firefly activity was then measured using a Tecan SPARK 10M. Renilla activity was measured after adding 30 μl of Dual-Glo Stop/Glo reagent.

Transactivation activity is expressed as the firefly/renilla RLU ratio. At least two independent experiments were performed with n=3-9.

For transactivation assays using primary CD4 T-cells, activated T-cells were transfected by the luciferase and the Tat vectors using the Amaxa program T23 as described by the manufacturer.

To assay histone acetylation, Jurkat cells were treated with drugs (D10, SAHA or BST-1) for 24 h before harvesting for SDS/PAGE and Western blots on nitrocellulose that were sequentially stained with anti-acetyl-histone (H2A, H2B, H3 or H4) and anti-histone (Cell Signaling Technology, kit #9933) before signal quantification using Fiji.

### LRA assay on cell lines

Cells (0.3.10^6^ /well) were plated in 24-well plates before adding drugs for 24 h. J-Lat 9.2 cells were washed once with PBS, then fixed with 3.7% Paraformaldehyde for 20 min at room temperature. Cells (>10,000) were then analyzed using a FACScalibur (Becton) or a Novocyte (Agilent) for the production of GFP that indicates HIV activation. Results are expressed as percentage of GFP positive cells. For ACH-2, J1.1 and OM10.1 cells, cells were centrifuged (3 min x 300 g) and the supernatant was treated with 1% Triton TX-100 to inactivate produced HIV-1 particles that were quantified using the INNOTEST HIV Antigen mAb kit (INNOGENETICS) p24 ELISA assay. Recombinant p24 was used as standard. At least three independent experiments were performed.

### LRA *ex vivo* assays

Cells (PBMCs or purified resting T-cells) were used immediately after purification. They were resuspended in RPMI containing 10% heat inactivated fetal calf serum at ∼10^6^ cells /ml and treated with drugs for 18 h in a 96-well plate. After centrifugation the cell supernatant was divided in two. Half was treated with 1% Triton TX-100 to inactivate HIV-1 particles that were quantified using a p24 ELISA (DuoSet ELISA, R&D Systems) using white 96-well plates and the protocol described by the manufacturer except for the final detection of HRP activity that was performed using Luminata Forte ELISA HRP substrate (Merk/Millipore). Using this modification, the detection limit is ∼0.1 pg p24 /ml. The second half was used to assay HIV RNA by qRT-PCR. RNA was first extracted using TRIzol LS using glycogen as a carrier for RNA precipitation. Reverse transcription was performed using All in one ABM Master mix, and qPCR was performed as described (Palmer *et al*, 2003), using Taqman universal master mix II (Applied Biosystem # 444040), a linearized Gag vector as standard, and a 384-well light cycler 480 (Roche).

### Mobility-shift electrophoresis assay

EMSA was performed using a procedure adapted from (Barboric *et al*., 2000). Recombinant GST-Tat (3 µM), P-TEFb (300 nM) or GST (3 µM) was incubated for 1h at room temperature with fluorescent TAR (250 nM) in 30 mM Tris pH 7.5 containing 100 mM KCl, 4 mM DTT, 1U RNAsin (Promega) / µl, 0.15% NP40, 2% glycerol, 2 mM MgCl_2_ and 10 µg polyIC (Invivogen) /ml. Samples were then loaded on 10% acrylamide gels that had been pre-run for 15 min at 4°C. Migration in 0.5x TBE was for 10 min at 30 V then 80 min at 100V and 4°C. Gels were then imaged for fluorescence using an Odyssey M gel scanner (Li-Cor). The resulting 32 bits images were analyzed using Fiji. Images shown in Fig.5 are contrasted to facilitate band viewing.

### Real-time PCR of cell-associated RNA

Cells were infected using VSV-G pseudotyped NL4.3 viruses (Schatz *et al*., 2023) using 20 and 200 ng p24 per million of Jurkat cells and CD4+ primary T-cells, respectively. After 18h, cells were washed 3 times with medium and resuspended in medium containing 5µM (for Jurkat cells) or 150 nM (for primary CD4 T-cells) D10 or SAHA. After 7h or 24 h, RNA was extracted and cDNA synthesized using TRIzol reagent (Invitrogen) and All-In-One Master Mix (ABM) kit as described by the manufacturers. Real-time PCR was performed using the LightCycler 480 SYBR Green I Master system (Roche). Standard curves were generated for all primer pairs using a serial dilution of linearized pNL4.3. Samples were amplified in triplicate in 5µL reactions containing 2.5µL 2X LightCycler SYBR Green I Master (Roche) and 2 µL of cDNA diluted 1:20. Data were analyzed using LightCycler 480 Software (Roche). PCR efficiencies ranged from 84% to 103%. Primers used were (Mousseau *et al*., 2012): GAPDH, 5’-AGGGATGACCTTGCCCACAGCCTTGG (Forward) and 5’ - CAACAGCCTCAAGATCATCAGCA (Reverse); HIV P3-NL4.3, 5’-GGCTAACTAGGGAACCCACTG (Forward) and P4, 5-CTGCTAGAGATTTTCCACACTGAC (Reverse), HIV P7, 5’-ACTTACGGGGATACTTGGGCAG (Forward) and P8 5’-CTCCATTTCTTGCTCTCCTCTGTC (Reverse), HIV P9, 5’ - GAAAAACATGGAGCAATCACAAGTAGCAATACAG (Forward) and P10, 5’ - CAGATCAAGGATATCTTGTCTTCTTTGGGAGTGAA (Reverse). These primers have been validated previously and enable the amplification of short transcripts (P3/P4; 42-158 bases of pNL4.3), medium transcripts (P7/P8; 5249-5358 bases), and long transcripts (P9/P10; 8444-8666 bases), respectively (Mousseau *et al*., 2012). HIV cDNA copy numbers were calculated using a digested pNL4.3 standard curve and the 2^-ΔΔCT^ method.

### pull-downs and immunoprecipitation

For GST-pull downs, HEK 293T cells were transfected with pGL3-LTR-luciferase using PEI Max (Schatz et al, 2023). After 18 h, D10 (5µM) or DMSO was added for 24 h. Cells were then harvested for preparation of nuclear extracts using a two-step protocol (Klenova *et al*, 2002). The extracts received 5 µM D10 when indicated. GST-Tat on glutathione-sepharose 4B beads (Cytiva) was then added before 2 h at 4°C on a rotating wheel, washes and resuspension in SDS/PAGE reducing sample buffer.

For D10-biotin pull down, HEK 293T cells were transfected with a Tat expression vector and pGL3-LTR-luciferase. D10-biotin or biotin (1 µM) were added to nuclear extracts before adding streptavidin magnetic beads (Dynabeads Myone streptavidin, Invitrogen # 65601), 16 h on a rotating wheel at 4°C, washes and resuspension in SDS PAGE reducing sample buffer.

For Tat immunoprecipitations, HEK 293 T cells were transfected with a Tat-FLAG expression vector with or without pGL3-LTR-luciferase. Nuclear extracts were then prepared, before adding anti-FLAG magnetic beads (Sigma M8823). After 90 min at 4°C on a rotating wheel, the resin was washed 4-fold in lysis buffer and resuspended in reducing sample buffer for SDS/PAGE. Western blots were stained for CDK9, CycT1 and Tat.

## List of Supplementary Materials

Fig S1 to S12

Table S1

## Acknowledgements

We are indebted to Bénedicte Fauvel and Henri Piras for discussions, Corinne Henriquet and Cyril Favard for help during SPR and fluorescence polarization experiments, respectively, to PLWHs that accepted to provide a blood sample for this study and to Camille Cregut for preliminary experiments. This work was funded by the Region Occitanie (Prematuration TatLat), the KIM "Biomarkers and Therapy" MUSE, the SATT AXLR (Grant 511-TATLAT) and Sidaction (grant # 20-2-AEQ-12908) to LC and BB. BB and LC hold a patent (WO2021233950) on the use of D10 as LRA. Author contribution. Conceptualization: BB, LC. Methodology: MP, JMP, LC, BB. Investigation: PBVT, LM, NC, MP, CB, LC, BB. Funding acquisition: BB, LC, AM, ET. Project administration and supervision: BB, LC, AM, ET. Writing – original draft: BB. Writing – review & editing: LC, BB, AM, ET. The authors declare that they have no competing interests.

---

**Figure S1.**
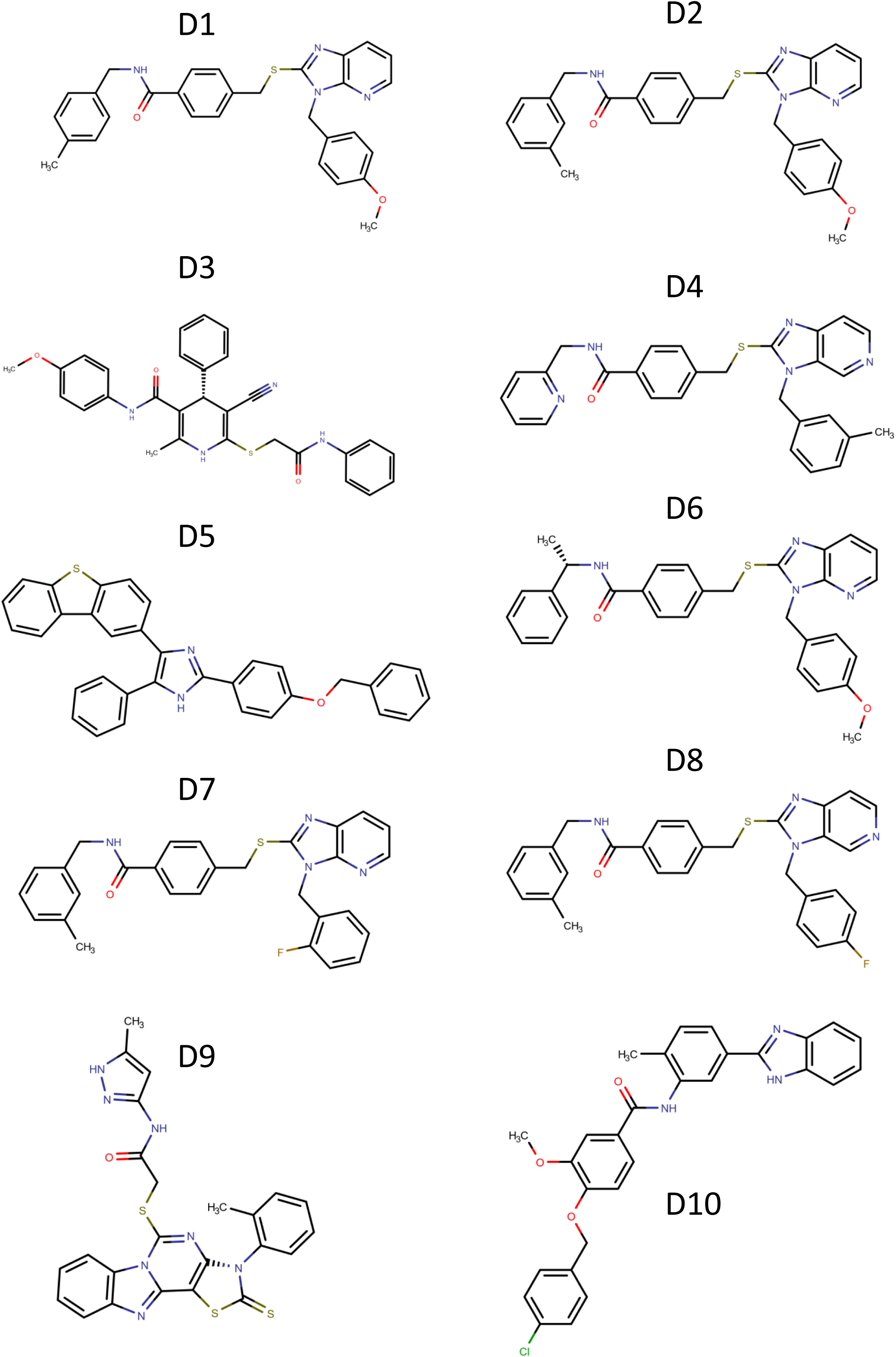
Structure of molecules from the D series.

**Figure S2.**
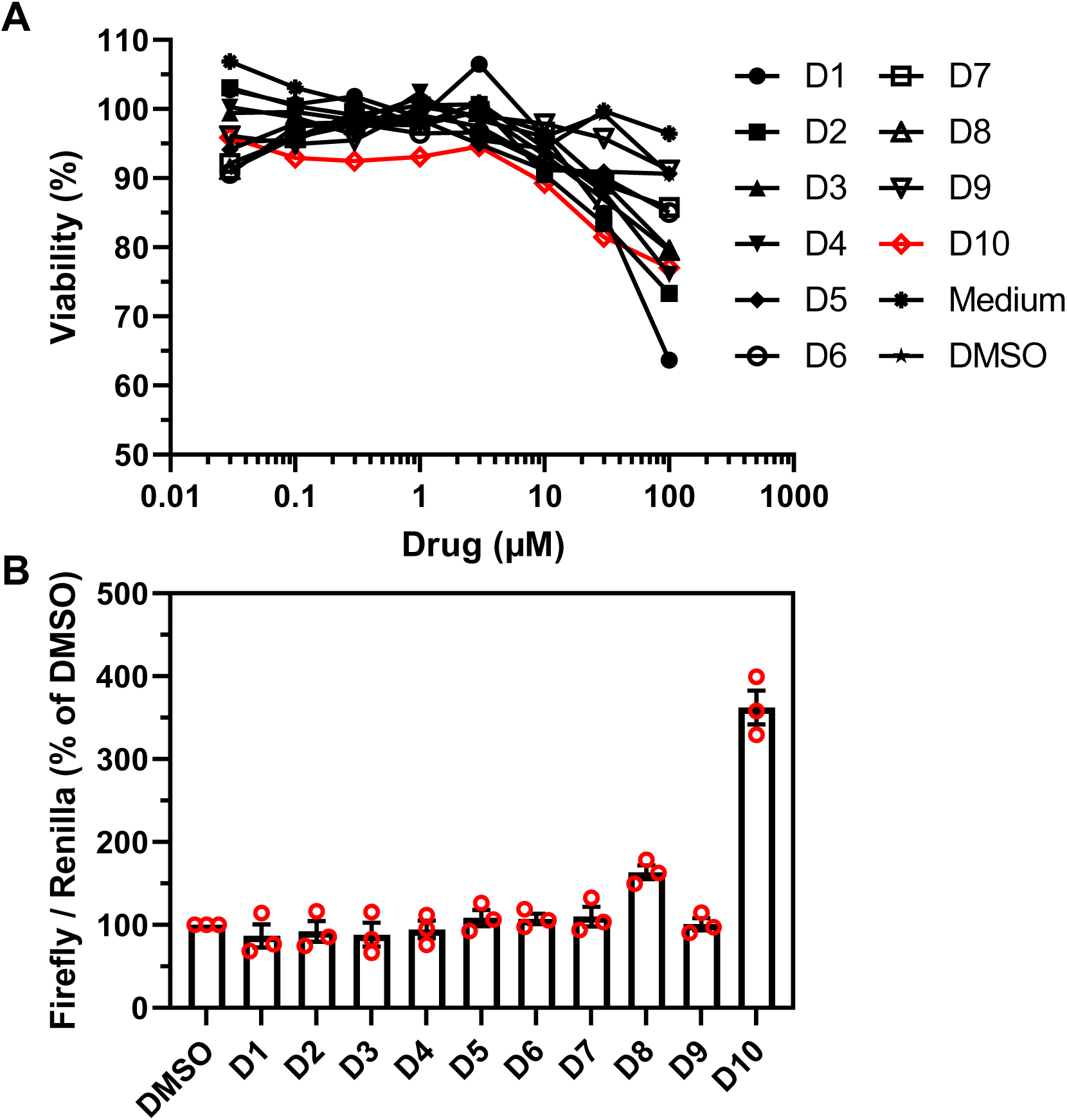
Cytotoxicity and transactivation assays of best Tat ligands. **A**, cytotoxicity assay. HeLa cells were treated with increasing doses of the molecules for 24 h, before assaying viability using Celltiter Blue. **B**, transactivation assay. HeLa cells were transfected with Tat and luciferase vectors before adding molecules at 5 µM for 24 h and luciferase assays. *Firefly* and *renilla* luciferases were behind HIV-1 LTR and a thymidine kinase promoter, respectively. Transactivation was followed using the Firefly / renilla activity ratio (AU). Data are mean ± SEM (n= 3 independent experiments).

**Figure S3.**
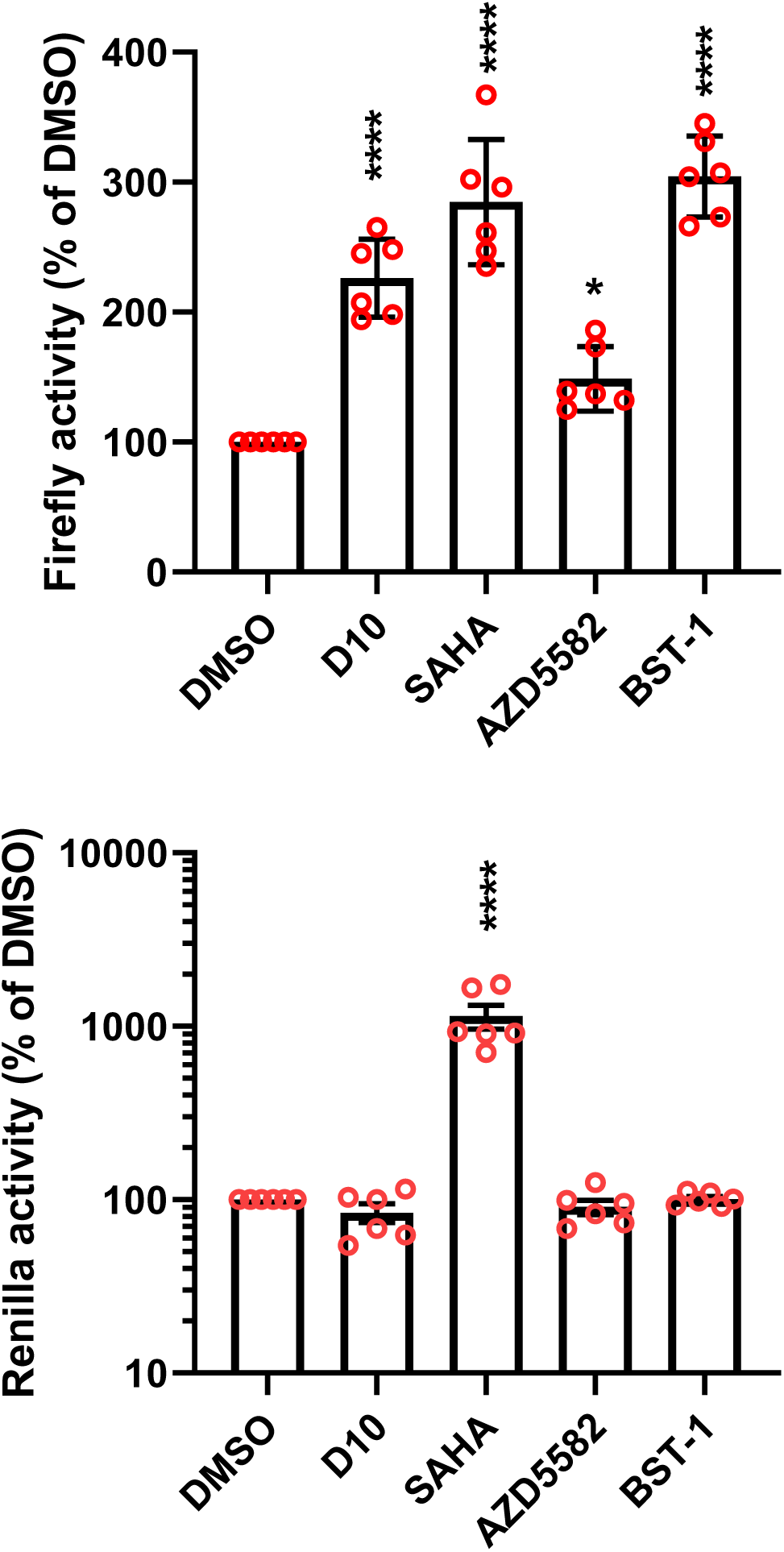
Transactivation activity of LRAs. HeLa cells were cotransfected with Tat, LTR-firefly and TK-renilla vectors. Drugs were added at 5 µM (except BST-1, 10 nM) after 18h. Cells were lysed for luciferase assays after a further 24h. Data are means ± SEM of n=3-7 independent experiments). One-way ANOVA. *, p< 0.05; ***, p<0.001; ****, p<0.0001.

**Figure S4.**
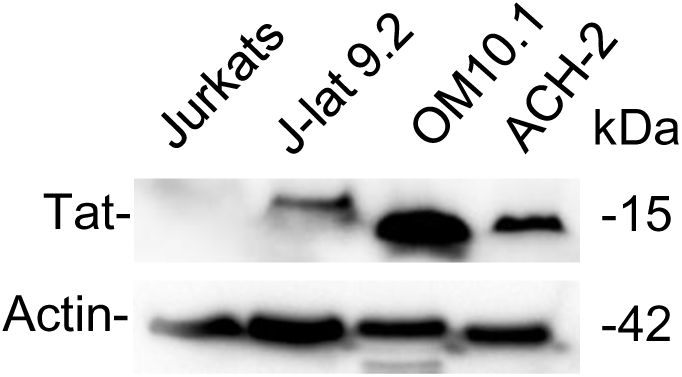
Tat is expressed in HIV latent cell lines. Cells (6 millions) were loaded on a 10 % Tricine gel before anti-Tat and anti-βactin Western blot.

**Figure S5.**
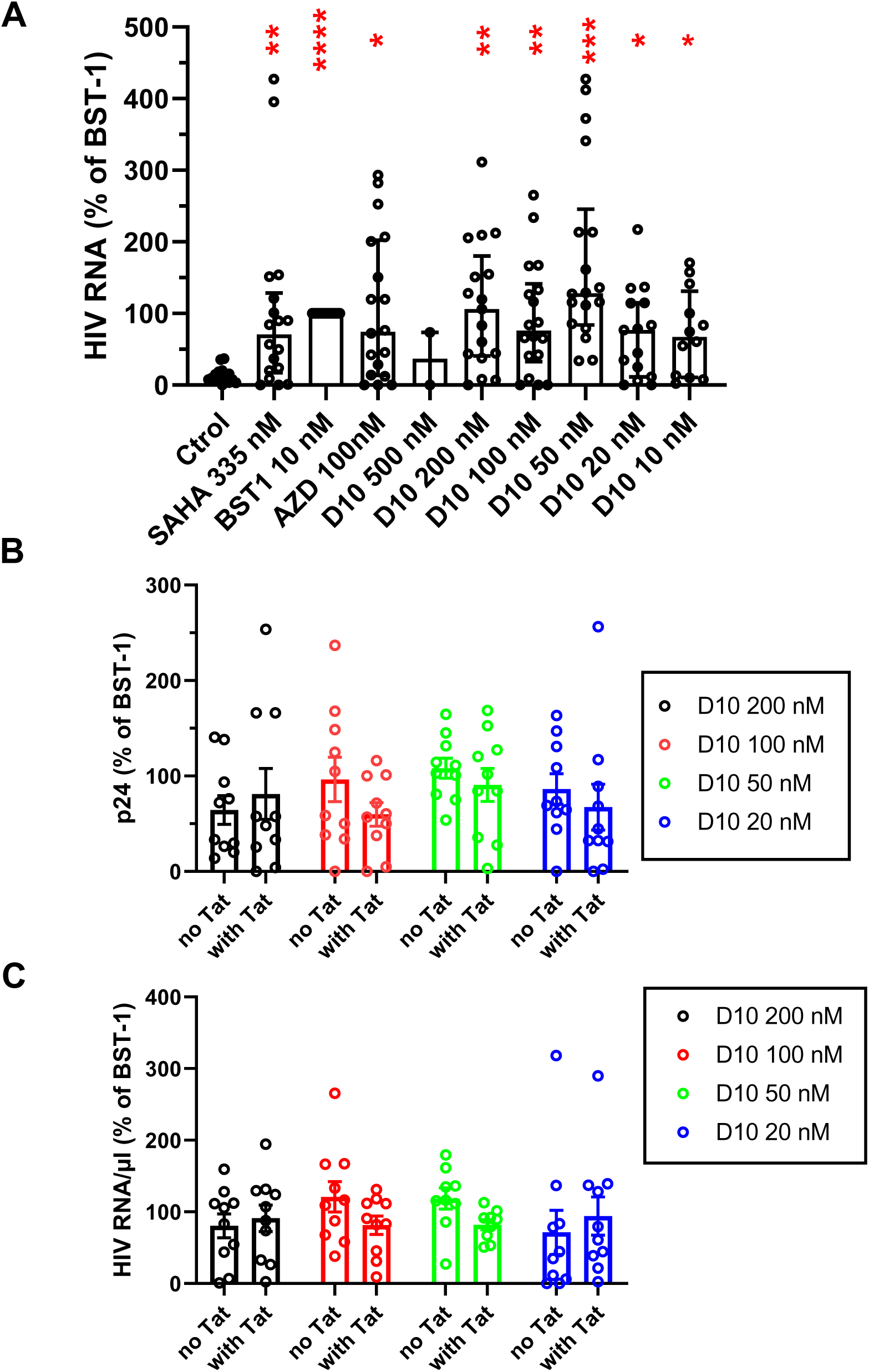
*Ex vivo* LRA activity. **A,** PBMCs were isolated from 18 PLWHs under ART. Cells in duplicated wells were incubated for 18-20 h in the presence of the indicated concentration of drug, before collecting supernatant for RNA extraction and qRT-PCR. Bryostatin-1 (BST1) response was set to 100% to enable easier comparison of LRA efficiency between PLWHs. Median ± interquartile range. Kruskal-Wallis tests. *, p<0.05; **, p<0.01; ***, p<0.001; ****, p<0.0001. **B** and **C,** The same experiment was performed in the presence of 10 nM recombinant Tat and viral production in the supernatant was followed using p24 ELISA (**B**) or qRT-PCR (**C**). No significant effect of exogenous Tat was observed (Two-Way ANOVA).

**Figure S6.**
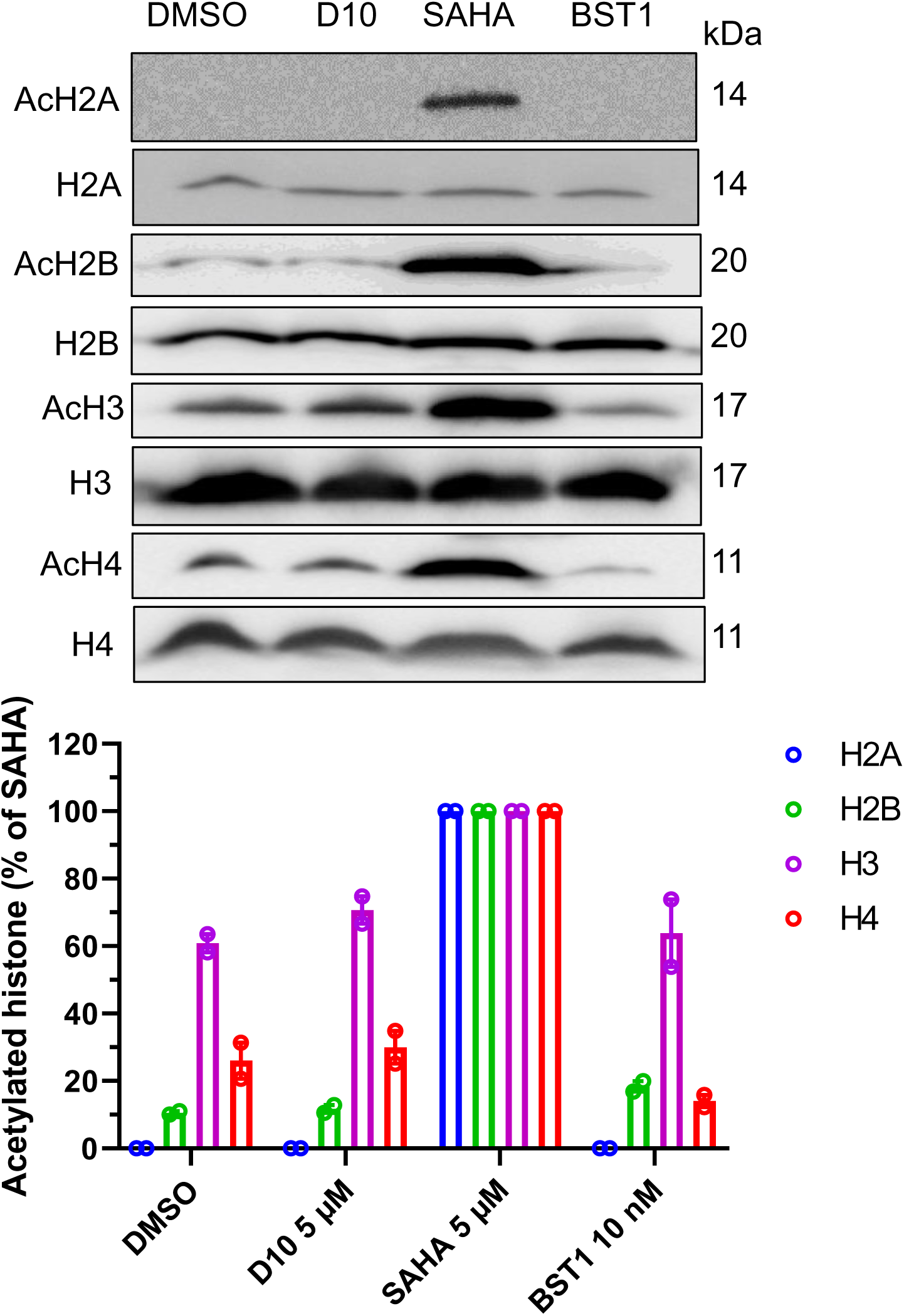
D10 is not an HDAC inhibitor. Jurkat cells were treated with the indicated drug concentration before lysis in SDS/PAGE sample buffer and histone acetylation analysis by western blots using specific antibodies. SAHA and bryostatin-1 (BST-1) were used as positive and negative controls, respectively. The graph presents quantification of the percentage of acetylated histone (mean ± SEM) from 2 independent experiment (each with n=2 blots). SAHA acetylation activity was set at 100% for each histone. No significant acetylation was induced by D10 or BST-1 compared to solvent (DMSO).

**Figure S7.**
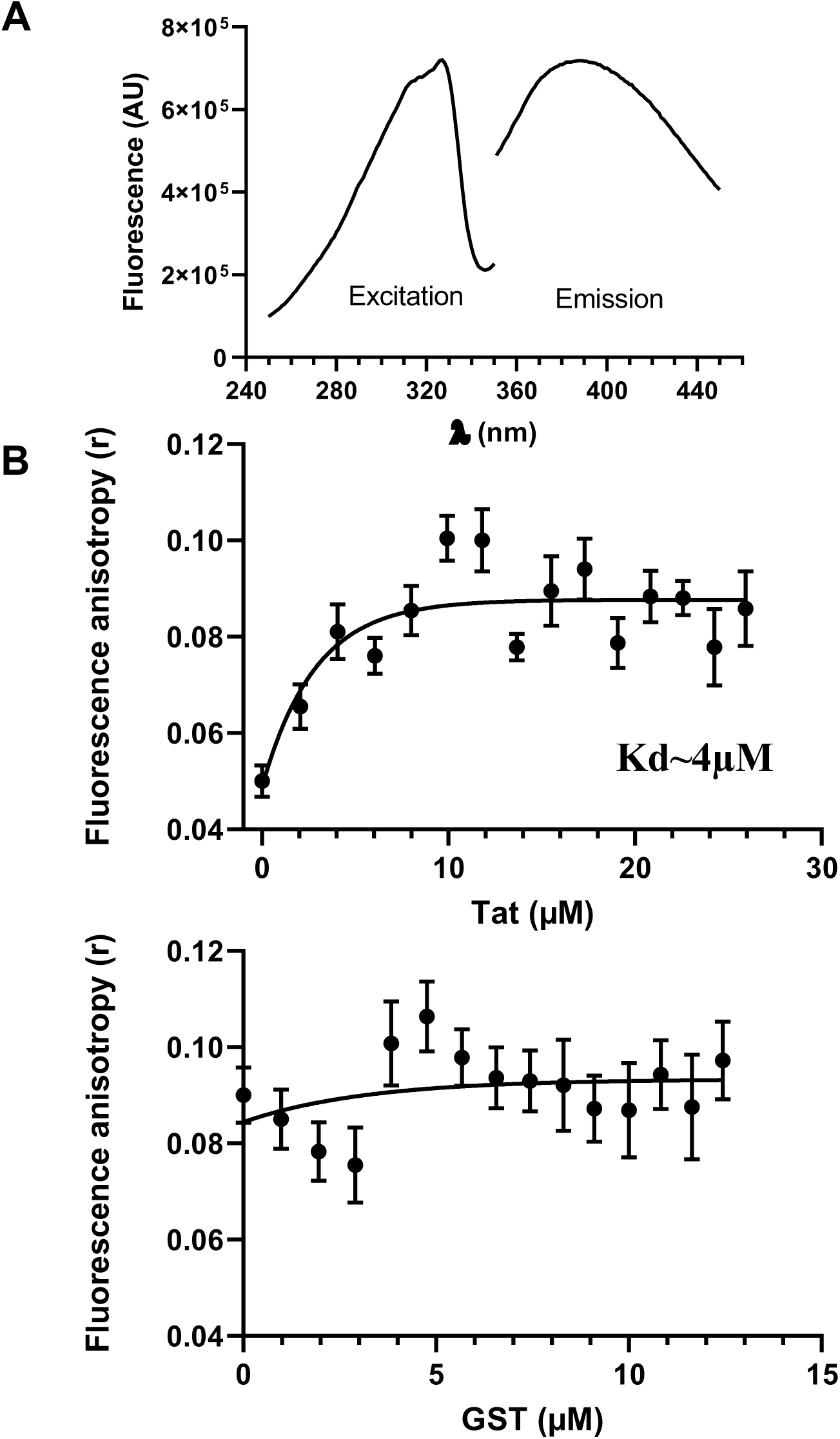
D10 fluorescence-polarization studies. **A,** Fluorescence spectra of D10. D10 was added at 1.25 µM in PBS and fluorescence spectra were recorded using a spectrofluorimeter. Solvent (DMSO) fluorescence represented ∼2% of D10 fluorescence and was subtracted. **B,** Fluorescence polarization of D10. D10 was added at 2 µM in a microcuvette containing citrate buffer and fluorescence polarization (at 0° and 90° with λ ex 320 nm and λ Em 410 nm) was recorded. Tat or GST were sequentially added from concentrated solutions (∼1 mM). Signals from solvent (DMSO) and proteins were subtracted and data were corrected from dilution. Bleaching was negligible over the experiment time frame. Anisotropy (r) was calculated using r=(I‖-I┴)/(I ‖+2I) (Rossi & Taylor, Nature Protocols 6 (2011) 365-387). Data are means ± SEM (n=3) and curves are exponential fits with one-phase decay.

**Figure S8.**
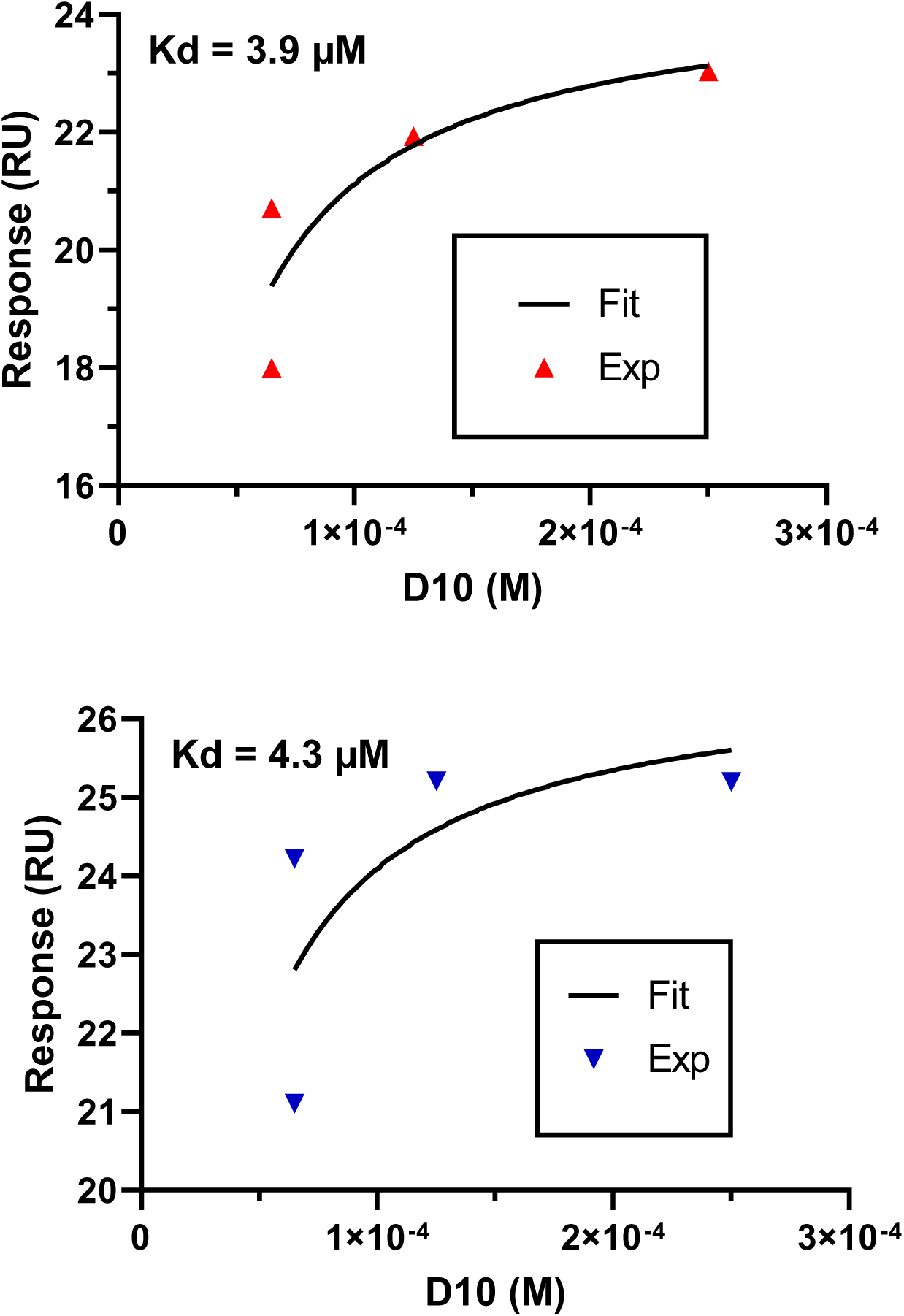
D10 binding to Tat followed using surface plasmon resonance. Tat was immobilized on a CM5 chip, before applying D10 (0-250 µM in citrate buffer). Two independent experiments are shown. Curves are exponential fits with one-phase decay for two independent experiments.

**Figure S9.**
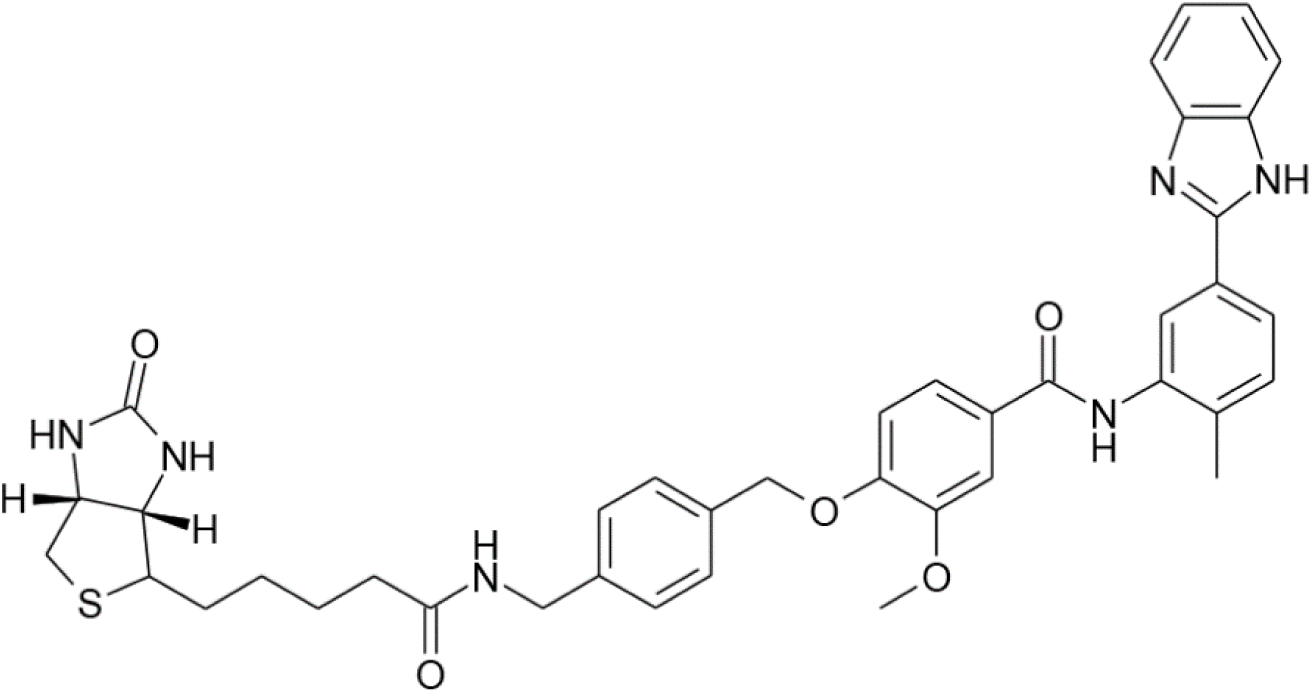
Structure of D10-biotin.

**Figure S10.**
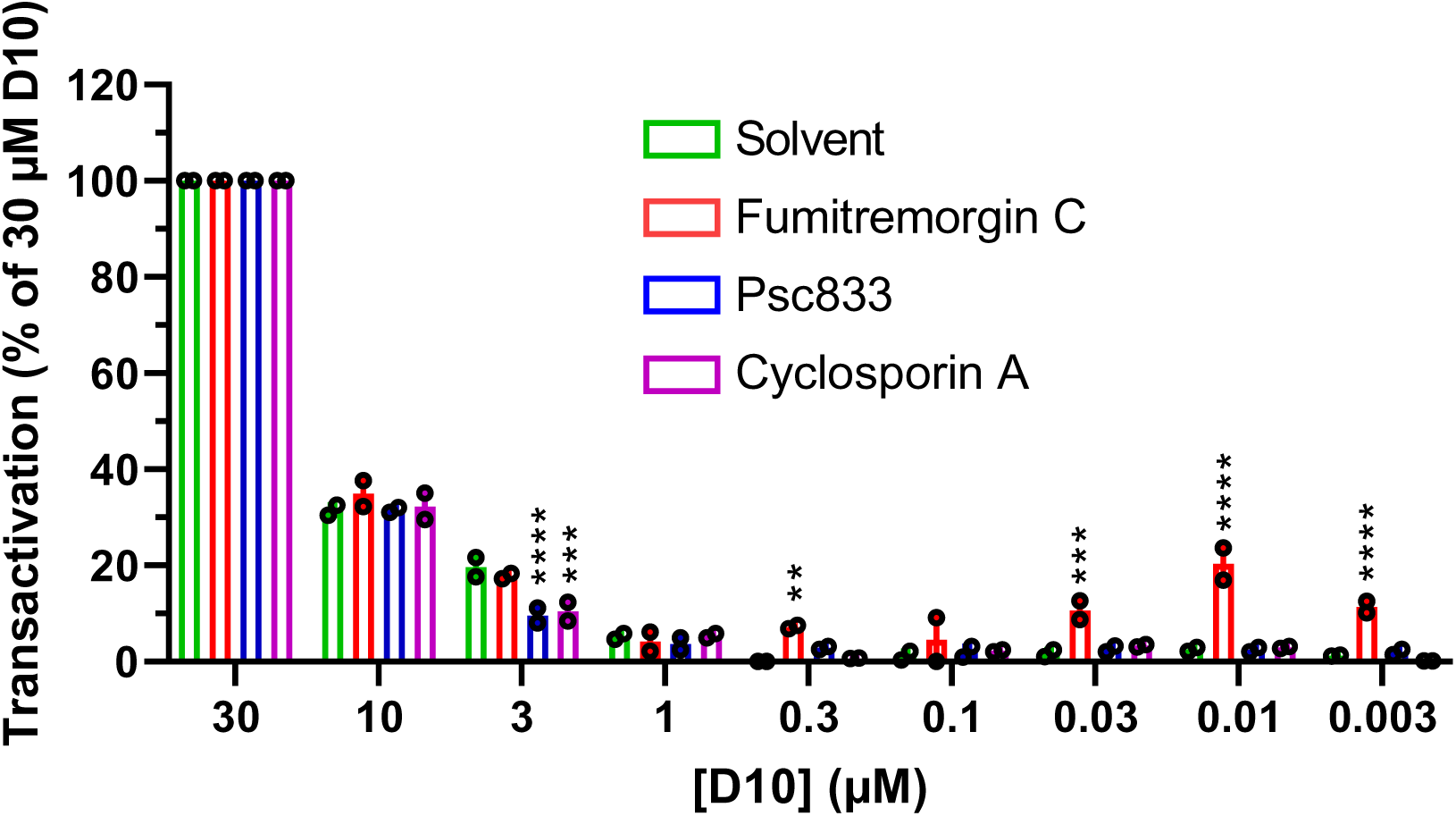
Inhibition of ABCG2 favors D10 effect on Tat transactivation in cell lines. HeLa cells were transfected with Tat and luciferase vectors before adding D10 together with the indicated inhibitor of ABCG2 (fumitremorgin C, 10 µM). Cyclosporin A (5 µM) and Psc833 (2 µM) that inhibit other transporters were used as controls. Cells were harvested after 24 h for luciferase assays. Transactivation was followed using the Firefly / renilla activity ratio. Data are mean ± SEM (n= 3 independent experiments). Two-way ANOVA compared to solvent (DMSO), **, p<0,01; ***, p<0,001; ****, p<0,0001.

**Figure S11.**
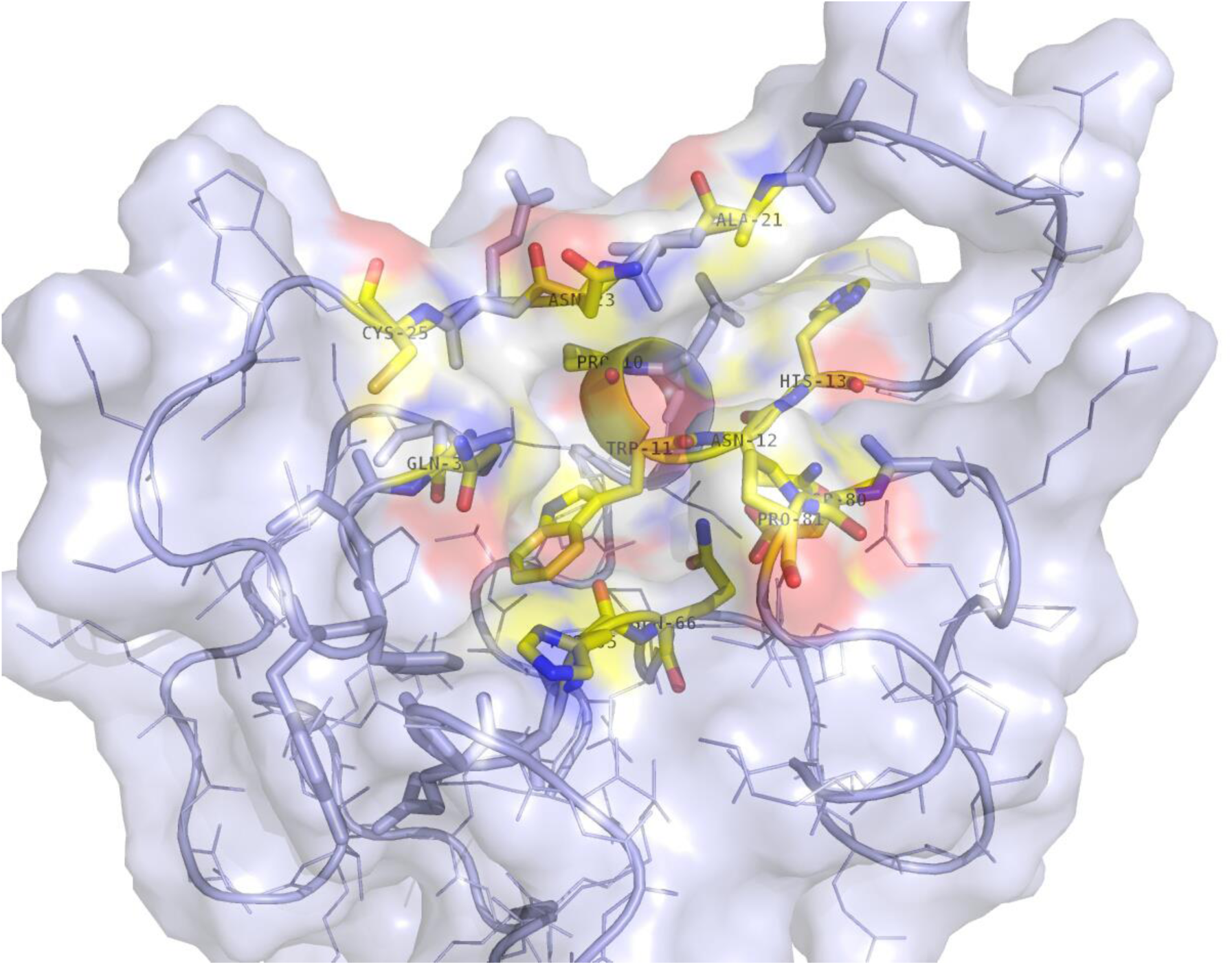
Conservation of D10 binding sites among HIV subtypes. 3D modeling was performed using subtype-D Tat. Residues involved in D10 binding appear in yellow. They are P10, W11, N12, H13, A21, N23, C25, Q35, H65, Q66, D80 and P81. In subtype-C, the Q35L mutation is present, while N12K and N23T are present in Tat subtype-B consensus sequence.

**Figure S12.**
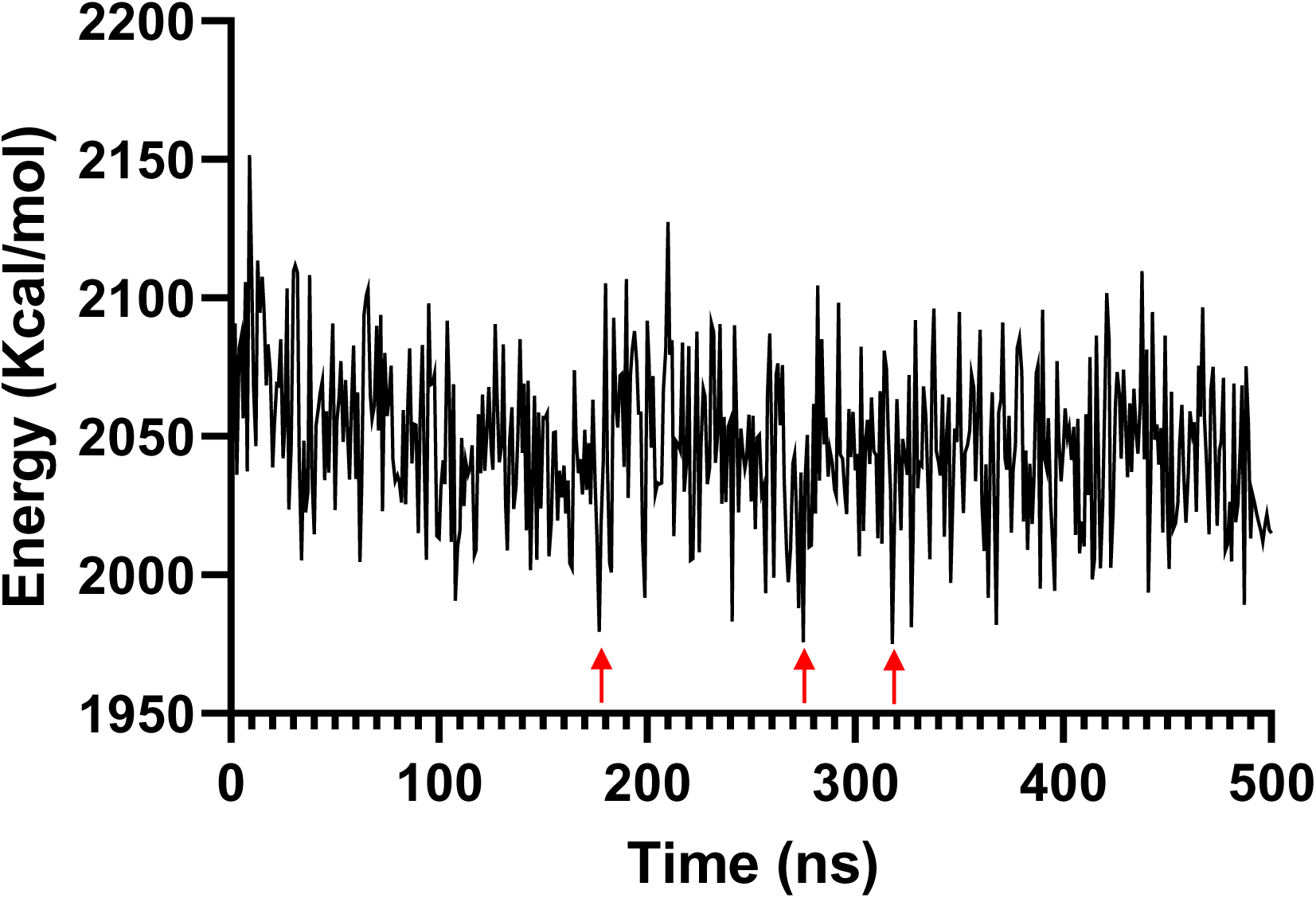
Conformational energy of Tat during 500 ns simulation. Conformational (non-bonded energy) component of the potential energy identifying the major stable conformations (*i.e.* conformers with lowest values of conformational energy) of Tat during 500 ns simulation. They are indicated by red arrows

**Table S1.**

| PLWH data for Fig.4A |  |  |  |  |  |  |  |  |  |  |
| --- | --- | --- | --- | --- | --- | --- | --- | --- | --- | --- |
| PLWH | Demography |  | Numeration |  | Infection HIV |  |  |  |  |  |
|  | SEX | Age | T-CD4 | T-CD8 | NADIR T-CD4 | Viral sub type | Indetectability | ART Composition | ART duration | Infection time |
|  |  |  | T4 / mm<sup>3</sup> | T8 / mm<sup>3</sup> | T4 / mm3 |  | months | Molecules | months | Years |
| 1 | M | 57 | 709 | 656 | 114 | B | 118 | JULUCA | 175 | 14 |
| 2 | M | 29 | 798 | 606 | 467 | CRF02_AG | 107 | VOCABRIA/REKAMBYS | 125 | 10 |
| 3 | M | 62 | 809 | 1252 | NK | B | NK | DELSTRIGO | 196 | 21 |
| 4 | M | 49 | 527 | 483 | NK | NK | 89 | BIKTARVY | 286 | 29 |
| 5 | F | 60 | 480 | 548 | 280 | NK | 246 | KIVEXA+VIRAMUNE | 369 | 34 |
| 6 | M | 35 | 739 | 449 | NK | B | NK | VOCABRIA/REKAMBYS | 116 | 10 |
| 7 | F | 53 | 580 | 378 | 190 | CRF18_cpx | 23 | TRUVADA/ISENTRESS | 73 | 6 |
| 8 | M | 78 | 495 | 323 | 215 | B | 12 | BIKTARVY | 156 | 13 |
| 9 | M | 76 | 980 | 501 | 361.85 | NK | 122 | DOVATO | 340 | 30 |
| 10 | M | 25 | 856 | 815 | 690 | CRF08_BC | 13 | BIKTARVY | 15 | 2 |
| 11 | M | 63 | 442 | 387 | 110 | B | 40 | DOVATO | 157 | 13 |
| 12 | M | 53 | 1189 | 443 | 361 | B | 174 | DOVATO | 178 | 15 |
| 13 | M | 52 | 653 | 788 | NK | B | NK | BIKARVY | NK | 28 |
| 14 | M | 55 | 403 | 543 | 6.88 | B | 94 | DOVATO | 388 | 40 |
| 15 | M | 47 | 604 | 653 | 238 | B | 120 | CABOTEGRAVIR/RILPIVIRINE | 128 | 11 |
| 16 | M | 63 | 321 | 596 | 50 | B | 141 | BIKTARVY | 334 | 35 |
| 17 | M | 50 | 1098 | 3567 | 497 | CRF02_AG | 104 | CABOTEGRAVIR/RILPIVIRINE | 129 | 10 |
| 18 | M | 69 | 794 | 285 | 794 | B | 109 | NORVIR/PREZISTA | 400 | 36 |

| PLWH data for Fig.4B |  |  |  |  |  |  |  |  |  |  |
| --- | --- | --- | --- | --- | --- | --- | --- | --- | --- | --- |
| 1 | M | 29 | 612 | 697 | NA | NA | 6 | NA | 6 | NA |
| 2 | M | 56 | 1208 | 1149 | NA | NA | 15 | NA | 15 | NA |
| 3 | M | 55 | 664 | 962 | NA | NA | 16 | NA | 21 | NA |
| 4 | M | 58 | 1358 | 1042 | NA | NA | 18 | NA | 21 | NA |
| 5 | F | 44 | 754 | 498 | NA | NA | 10 | NA | 21 | NA |
| 6 | M | 61 | 997 | 457 | NA | NA | 11 | NA | 11 | NA |
| 7 | F | 66 | 1282 | 1069 | NA | NA | 3 | NA | 3 | NA |
| 8 | M | 50 | 758 | 961 | NA | NA | 4 | NA | NA | NA |
| 9 | M | 47 | 1008 | 1607 | NA | NA | 20 | NA | NA | NA |

| Inclusion criteria | IC_1 | Age > 18 y |
| --- | --- | --- |
|  | IC_2 | People living with HIV |
|  | IC_3 | vRNA undetectable for >12 months |
| Non inclusion criteria | NIC_1 | Absence of ART treatment |
|  | NIC_2 | ART just beginning |
|  | NIC_3 | immunosuppressive medication |
|  | NIC_4 | Cancer less than 5 years ago |
|  | NIC_5 | Pregnant or breastfeeding woman |
|  | NIC_6 | Law protected person |
|  | NIC_7 | People involved in another research project |
|  | NIC_8 | People not benefiting from French social security system |
|  | NIC_9 | People refusing to participate in this study |

NA not available

